# Somatostatin interneurons control the timing of developmental desynchronization in cortical networks

**DOI:** 10.1101/2023.12.21.572767

**Authors:** Laura Mòdol, Monika Moissidis, Oscar Marín

**Author notes:** Correspondence (LM), (OM).

## Abstract

Synchronous neuronal activity is a hallmark of the early developing brain. In the mouse cerebral cortex, activity decorrelates during the second week of postnatal development, progressively acquiring the characteristic pattern of sparse coding underlying the integration of multidimensional sensory information. The maturation of inhibition seems critical for this process, but the specific types of interneurons involved in this crucial transition of network activity in the developing cortex remain unknown. Using in vivo volumetric and longitudinal two-photon calcium imaging during the period that precedes the change from highly synchronous to decorrelated activity, we identify somatostatin-expressing (SST+) interneurons as critical modulators of this switch. Modulation of the activity of SST+ cells accelerates or delays the decorrelation of cortical network activity, a process that involves regulating the degree of maturation of parvalbumin-expressing (PV+) interneurons. SST+ cells critically link sensory inputs with local circuits controlling the neural dynamics in the developing cortex while modulating the integration of other interneurons into nascent cortical circuits.

---

During early postnatal development, cortical networks are characterized by oscillatory bursts of activity synchronizing large numbers of neurons^1–4^. These synchronous activity patterns decorrelate over time, reflecting changes in the connectivity and maturation of emerging neural circuits^5–7^. The decorrelation of spontaneous activity leads to sparse coding, a ubiquitous strategy critical for information processing across multiple cortical areas^8^. Although the transition of neural activity from highly synchronous to sparse and decorrelated is a crucial milestone in the development of functional cortical networks, the cellular mechanisms underlying this process remain poorly understood.

Theoretical and experimental studies indicate that the balance between excitation and inhibition is critical for sparse coding^9–11^. During early postnatal development, the ratio of excitatory and inhibitory conductances in cortical neurons is higher than in the adult brain. However, this ratio decreases as cortical networks mature due to a progressive increase in inhibition^12^, suggesting that the functional integration of GABAergic interneurons into emerging neural circuits is required to sparsify activity in the cerebral cortex. Consistent with this notion, experimental manipulation of cortical interneurons during early postnatal development influences the spatiotemporal structure of spontaneous activity patterns^13–17^ and the decorrelation of cortical activity^18^.

The adult cerebral cortex contains a vast diversity of GABAergic interneurons with unique electrophysiological properties and connectivity, which endows inhibition with a wide range of mechanisms to shape cortical computations^19–21^. Interneurons develop over a protracted period before they acquire their mature features. Notably, different types of interneurons follow distinct developmental trajectories, integrating into cortical circuits asynchronically^22^. For example, interneurons derived from the medial ganglionic eminence (MGE), which comprise two main subclasses, somatostatin-expressing (SST+) interneurons and parvalbumin-expressing (PV+) interneurons, integrate into cortical circuits before interneurons derived from the caudal ganglionic eminence^22,23^. Among the MGE-derived interneurons, SST+ cells mature faster than PV+ interneurons^24–26^ and influence cortical dynamics during the first week of postnatal development^27–29^. Thus, it is likely that specific types of interneurons regulate the transition from highly synchronous activity to sparse coding in developing cortical networks.

Here, we combined longitudinal two-photon calcium imaging in vivo and pharmacogenetics to investigate the role of SST+ and PV+ interneurons in modulating network activity during the first two weeks of postnatal development in the mouse. We found that SST+ interneurons are critical for regulating spontaneous and sensory-evoked synchronous activity patterns in the barrel cortex, promoting the synchronization of cortical activity during the first week of postnatal development. Inhibiting SST+ interneurons during this period accelerates the maturation of PV+ interneurons and the decorrelation of cortical activity, suggesting that SST+ cells modulate cortical dynamics at least partially by regulating the integration of other interneurons into nascent cortical circuits. Our results indicate that the sequential integration of SST+ and PV+ interneurons into cortical circuits determines the transition of cortical activity from highly synchronous to sparse and decorrelated during the second week of postnatal development.

## RESULTS

### Spatiotemporal decorrelation of spontaneous activity in the developing cortex

To understand the mechanisms driving the decorrelation of neural activity in the mouse neocortex during early postnatal development, we first investigated the neuronal dynamics underlying this developmental switch. To this end, we performed longitudinal two-photon calcium imaging in pups from postnatal day (P)7 to P12, when cortical neuronal activity begins to sparsify^5,6^. We injected the barrel field of the primary somatosensory cortex (S1BF) of P0 pups with a pan-neuronal adeno-associated virus (AAV) encoding the calcium sensor GCaMP6s to examine the activity of a large population of cortical neurons (Figure 1A). We then monitored changes in spontaneous activity during the second week of postnatal development in vivo by imaging a field of 650 x 650 µm at different cortical depths and up to 500 µm below the surface in S1BF on consecutive days (Figure 1B). We used a deep-learning-based tool (CASCADE) to estimate neuronal firing rates from changes in the fluorescent signal of calcium imaging^30^. Calcium events from the resulting binarized matrices of spikes were then detected using a surrogate distribution (Figure S1A). Individual imaging sessions were pooled within two consecutive days to obtain a dataset comprising three developmental stages (P7/8, P9/10, and P11/12) that captured the dynamic changes of spontaneous activity in S1BF during early postnatal development.

**Figure 1.**
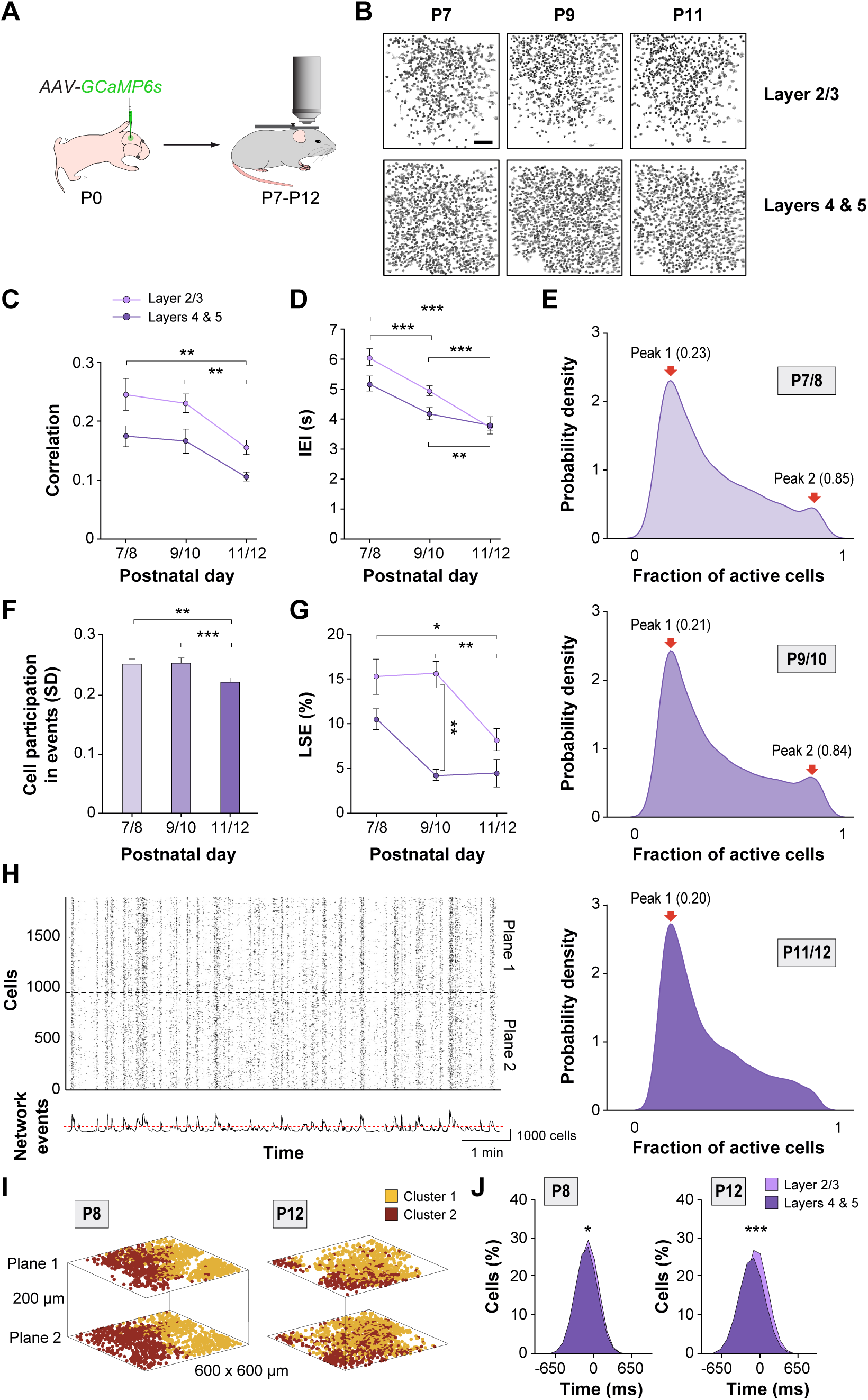
Spatiotemporal decorrelation of spontaneous activity during postnatal development. (A) Schematic representation of the experimental paradigm. Imaging was performed in superficial (L2/3: 100-250 µm depth; 97 field of views (FOVs) from 7 pups) and deep (L4/5: 300-500 µm depth; 33 FOVs from 5 pups) layers of pups between P7 and P12. (B) Contour map of active cells in L2/3 and L4/5 from a representative example following the same field of view (FOV) at P7, P9, and P11. (C) Developmental changes in the correlation degree of active cells in L2/3 and L4/5. Two-way ANOVA for age: F(2,124) = 6.85, *p* < 0.001. Post hoc Tukey’s multiple comparison test indicates differences in L2/3 between P7/8 and P11/12 (\*\**p <* 0.01), and P9/10 and P11/12 (\*\**p* < 0.01). No differences were observed between L2/3 and L4/5 at any stage. (D) Developmental changes in Inter Event Intervals (IEI) duration in L2/3 and L4, and L5. Two-way ANOVA for age: F(2,124) = 28.84*, p* < 0.0001. Post hoc Tukey’s multiple comparison test indicates differences in L2/3 between P7/8 and P9/10 (\*\*\**p* < 0.001), P7/8 and P11/12 (\*\*\**p* < 0.001) and P9/10 and P11/12 (\*\*\**p* < 0.001) and in L4 and L5 between P7/8 and P11/12 (\*\**p* = 0.01). No differences were observed between L2/3 and L4/5 at any stage. (E) Probability density estimation of cells recruited within spontaneous events at different developmental stages (P7/8, P9/10, and P11/12). The peaks indicate the probability density of cell recruitment, whereas the numbers in parenthesis indicate the fraction of cells recruited at the peak. (F) Differences in the standard deviation of cell recruitment within spontaneous calcium events at different postnatal stages. One-way ANOVA: F(2,124) = 2.38*, p* < 0.001. Post hoc Tukey’s multiple comparison test indicates differences between P7/8 and P11/12 (*\*\*p* < 0.01) and between P9/10 and P11/12 (\*\*\**p* = 0.001). (G) Developmental changes in the fraction of Large Synchronous Events (LSEs) in L2/3 and L4/5. Two-way ANOVA for age F(2,124) = 5.52, *p* < 0.01. Two-way ANOVA for depth F(1,124) = 14.84, *p* < 0.01. Post hoc Tukey’s multiple comparison test indicates differences in L2/3 between P7/8 and P11/12 (\**p* < 0.05) and P9/10 and P11/12 (\*\**p* < 0.01). (H) Raster plots illustrating calcium events during simultaneous imaging of L2/3 and L4/5 at P8. Plane 1 corresponds to the most superficial imaged plane located at 150 µm in depth (L2/3). Plane 2 corresponds to the deepest imaged plane at 350 µm (L4/5). The cumulative histogram of network events indicates the sum of active contours in the raster plots over time. The red dotted line indicates the sum of spikes considered a network event based on a statistical threshold (*p* < 0.01). (I) Spatial correlation between cells during simultaneous imaging of L2/3 and L4/5 at P8 (6 FOVs from 4 pups) and P12 (3 FOVs from 3 pups). The colors indicate cells corresponding to the same or different assembly (k-means clustering) along the x, y, and z-axis. (J) Quantification of cell recruitment in spontaneous events in L2/3 (light purple) and L4/5 (dark purple) at P8 and P12. In the x-axis, 0 corresponds to the peak of cell recruitment in events. Each distribution represents the fraction of participation in spontaneous events scaled to the maximum participation rate. *t*-tests: \**p* < 0.05 for P8 and \*\*\**p <* 0.001 for P12. Data is presented as mean ± s.e.m. Scale bar, 100 µm (B).

Consistent with previous studies^5^, we found that spontaneous activity in S1BF gradually changes from highly correlated to uncorrelated during the second postnatal week of the mouse’s life (Figure 1C). We also noticed that the sparsification of spontaneous activity is accompanied by an increase in the frequency of events (i.e., a decrease in Inter Event Intervals, IEIs) across both superficial and deep cortical layers (Figure 1D). Moreover, the analysis of the number of active cells within events revealed an important milestone accompanying the functional decorrelation of spontaneous activity. We observed that the participation of neurons in network events at P7/8 and P9/10 is characterized by a binomial distribution, with network events recruiting about 20% or, less frequently, more than 85% of the cells (Figures 1E and S1). This pattern changes to a unimodal distribution at P11/12 when the large synchronous events (LSEs) involving more than 85% of the active cells become less frequent (Figures 1E and S1B). A comparison of the standard deviation of cell participation in events further confirmed a significant decrease in the variability of event types at P11/12 (Figure 1F). In other words, during the decorrelation of network dynamics, cortical activity becomes more recurrent, but network events involve fewer cells than at earlier stages. Analysis of the prevalence of LSEs across cortical layers revealed that this pattern of activity disappears earlier in granular and infragranular cortical layers than in supragranular layers (Figure 1G), which is generally consistent with the progressive maturation of cortical connectivity following an inside-out pattern^31,32^.

We analyzed the spatiotemporal properties of spontaneous activity across cortical layers and developmental stages in greater detail. To this end, we performed volumetric two-photon in vivo calcium imaging of S1BF from P7 to P12, which allows the analysis of spontaneous activity simultaneously recorded across multiple layers. Consistent with previous electrophysiological studies in slices^25,33^, we observed that calcium events span through the cortical depth and recruit cells across different layers at the beginning of the second postnatal week (Figure 1H). K-means clustering of activity correlations revealed that cells at different depths form large functional assemblies in spatially restricted columns as early as P7/8. This functional columnar organization disappears at P11/12 when the activity starts to decorrelate (Figure 1I and S1C). We also noticed that cells located in granular and infragranular layers of the cortex fire earlier than cells located in supragranular layers at P7/8 and P11/12, although this temporal pattern becomes much more distinctive at the later stages (Figure 1J). This temporal pattern reflects the role of granular and infragranular layers in directing cortical dynamics. Overall, our results indicate that the decorrelation of spontaneous activity is accompanied by significant changes in the functional connectivity of superficial and deep layers of the neocortex, with deep layers likely driving the generation of spontaneous events.

### Functional integration of SST+ and PV+ interneurons in cortical network activity

Previous work has suggested that a relative increase in inhibition mediates the decorrelation of spontaneous activity in the developing prefrontal cortex^18^. To explore the role of distinct interneurons in the sparsification of cortical activity, we focused on SST+ and PV+ interneurons, two subclasses of cortical interneurons that populate granular and infragranular layers from early stages of postnatal cortical development^34,35^. To this end, we crossed *Sst^Cre/+^* and *Tac1^Cre/+^* mice with *RCL^tdTomato/+^* (Ai9) reporter mice to label SST+ and PV+ interneurons, respectively. In separate experiments, we injected the S1BF of *Sst^Cre/+^;Ai9* and *Tac1^Cre/+^;Ai9* pups at P0 with a cocktail of pan-neuronal and Cre-dependent AAVs expressing GCaMP6s and monitored the activity of cortical cells from P7 to P12 in vivo (Figure 2A). Because SST is expressed in migrating interneurons embryonically, *Sst^Cre/+^;Ai9* mice identify a significant fraction of SST+ interneurons in the early postnatal cortex^36^. Conversely, PV only begins to be expressed in S1BF from P10 onwards^37^, but *Tac1^Cre/+^;Ai9* mice allow the identification of a subset of prospective PV+ interneurons before the onset of PV expression^38^. Using this strategy, we monitored the activity of a relatively large (>40%) sample of SST+ and PV+ interneurons in the early postnatal cortex along with many other cells, which broadly correspond to pyramidal cells (Figures 2B and S2A–S2C).

**Figure 2.**
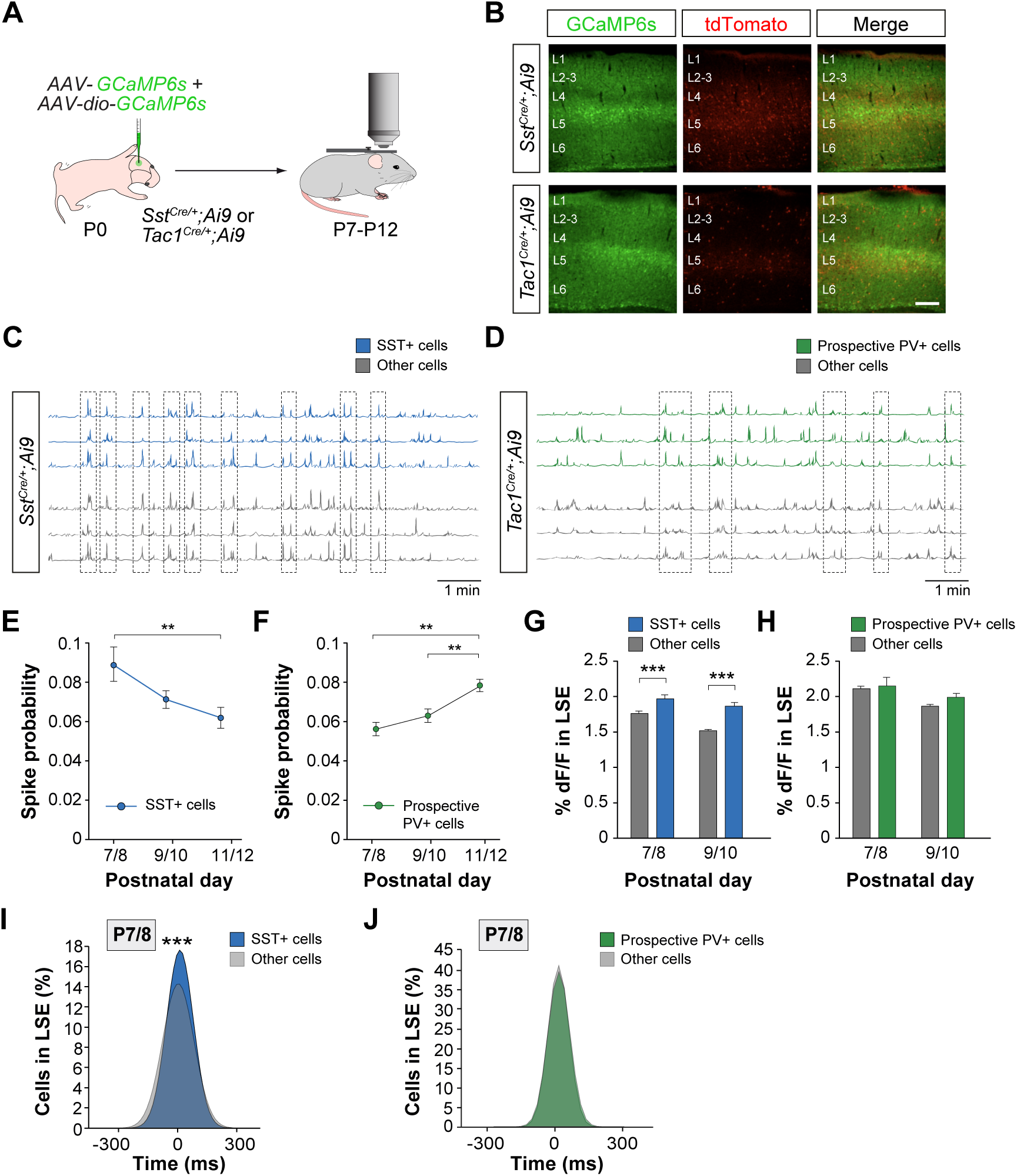
Divergent integration of SST+ and prospective PV+ interneurons in developmental activity patterns. (A) Schematic representation of the experimental paradigm. *Sst^Cre/+^;Ai9* mice: 9 pups (P7/8), 10 pups (P9/10), and 9 pups (P11/12). *Tac1^Cre/+^;Ai9* mice: 3 pups (P9/10), 6 pups (P9/10), and 6 pups (P11/12). (B) Coronal sections illustrating the expression of GCaMP6s and tdTomato in the S1BF of *Sst^Cre/+^;Ai9* and *Tac1^Cre/+^;Ai9* at P7/8. (C) Raster plots illustrating calcium traces from SST+ (blue) and other cells (grey). (D) Raster plots illustrating calcium traces from Tac1+ (green) and other cells (grey). (E and F) Changes in spike probability of SST+ and Tac1+ cells across developmental stages. One-way ANOVA for each cell type; SST+: F(2,25) = 4.59*, p* = 0.01. Post hoc Tukey’s multiple comparison test indicates differences in the firing probability between P7/8 and P11/12 (\*\**p* < 0.01). Tac1+: F(2,12) = 10.16*, p* < 0.01. Post hoc Tukey’s multiple comparison test indicates differences in the firing probability between P7/8 and P11/12 (\*\**p* < 0.01) and between P9/10 and P11/12 (\*\**p =* 0.01). (G) Quantification of calcium traces (dF/F) during Large Synchronous Events (LSEs) for SST+ cells (blue) and other active cells (grey) in *Sst^Cre/+^;Ai9* as a function of age. Two-way ANOVA for age and cell type: Interaction effect: F(1,55996) = 4.94*, p* < 0.05. Cell type effect: F(1,55996) = 33.87*, p* < 0.0001 and Age effect: F(1,55996) = 84.88*, p* < 0.0001. Post hoc Tukey’s multiple comparison test indicates differences between SST+ cells and other active cells at P7/8 (\*\*\**p* < 0.0001), P9/10 (\*\*\**p <* 0.0001). (H) Quantification of calcium traces (dF/F) during Large Synchronous Events (LSEs) for Tac1+ cells (green) and other active cells (grey) in *Tac1^Cre/+^;Ai9* as a function of age. Two-way ANOVA for age and cell type: Interaction: F(1,17728) = 0.70*, p* = 0.40. Cell type: F(1,17728) = 1.61*, p* = 0.20. Age: Cell type: F(1,17728) = 16.76*, p* < 00001. (I) Quantification of cell participation in Large Synchronous Events (LSEs) for SST+ cells (blue) and other active cells (grey) in *Sst^Cre/+^;Ai9* as a function of age. Shadowed distributions illustrate the probability of cells participating in LSEs scaled to the maximum participation rate. *t*-test: \*\*\**p* < 0.0001. (J) Quantification of cell participation in Large Synchronous Events (LSEs) for Tac1+ cells (green) and other active cells (grey) in *Tac1^Cre/+^;Ai9* as a function of age. Shadowed distributions illustrate the probability of cells participating in LSEs scaled to the maximum participation rate. *t*-test: *p* = 0.56. Data is presented as mean ± s.e.m. Scale bar, 100 µm (B).

We observed that SST+ and prospective PV+ interneurons fire concomitantly with the rest of the cortical neurons, integrating into the spontaneous activity patterns that characterized cortical activity between P7 and P12 (Figures 2C, 2D, and S2B-S2D). Next, we investigated the spiking probability of SST+ and prospective PV+ interneurons. We observed that SST+ and prospective PV+ cells display an inverse activation pattern in S1BF during the second week of postnatal development. SST+ are significantly more active at the beginning of the second postnatal week (P7/8) when activity is highly correlated and cortical activity is characterized by LSEs (Figure 2E). In contrast, the spiking probability of putative PV+ cells is relatively low at P7/8 but increases progressively over the next few days (Figure 2F). These observations suggested that SST+ interneurons may have a particularly critical role in regulating cortical activity at the beginning of the second postnatal week when prospective PV+ interneurons are still relatively inactive.

To further investigate this question, we analyzed the contribution of SST+ and prospective PV+ cells to specific network events. To this end, we examined the amplitude of calcium transients (% dF/F) in SST+ and putative PV+ cells during LSEs. We found that SST+ cells but not prospective PV+ interneurons display larger calcium responses than other neurons at the beginning of the second postnatal week (Figures 2G and 2H), when LSEs are particularly abundant in S1BF (Figures 1E–1G). We also observed that SST+ cells participating in LSEs fire significantly later than other neurons (putative pyramidal cells) at P7/8 (Figure 2I). This activation pattern was exclusively found during LSEs and not in general network events (Figure S2E), reinforcing the notion that SST+ interneurons function as a break of run-away excitation during this specific developmental period characterized by frequent LSEs^39^. Interestingly, we observed that the pattern of temporal correlations between prospective PV+ interneurons and other neurons (putative pyramidal cells) firing in events was reversed to that found for SST+ interneurons, with prospective PV+ interneurons firing significantly later than pyramidal cells only in events at P11/12 (Figures 2J and S2F). This observation illustrates the gradual integration of PV+ interneurons into functional circuits with pyramidal cells, which is significantly delayed compared to SST+ interneurons.

### Silencing SST+ interneurons restrict neuronal participation in LSEs

Our previous results revealed that the integration of SST+ and prospective PV+ interneurons into functional circuits in S1BF is temporally shifted during the second week of postnatal development. These observations also suggested that SST+ interneurons may be critical in regulating cortical network activity before PV+ cells become functionally mature. To test this idea, we suppressed the activity of cortical SST+ cells using a chemogenetic approach based on Designer Receptors Exclusively Activated by Designer Drugs (DREADDs)^40^. In brief, we injected S1BF of P0 *Sst^Cre/+^;Ai9* pups with a cocktail of pan-neuronal and Cre-dependent AAVs expressing GCaMP6s and Cre-dependent AAVs encoding mutant G-protein-coupled receptors that induce neuronal inhibition (hM4Di) following administration of the pharmacologically inert molecule clozapine-*N*-oxide (CNO). We then injected P8 mice with CNO or vehicle and monitored neuronal activity 10 min later using two-photon calcium imaging in vivo (Figure 3A). We observed that reducing the activity of SST+ cells by CNO administration (Figures S3A–S3D) increased the frequency of network events (Figures 3B and 3C). We also found that inhibition of SST+ interneurons promotes the generation of events that display shorter duration and recruit fewer cells than in control mice (Figures 3D and 3E). Consistently, the fraction of LSEs in CNO-injected mice was significantly lower than in control mice (Figure 3F). These observations indicate that inhibition of SST+ cells promotes the decorrelation of spontaneous activity at the beginning of the second week of postnatal development.

**Figure 3.**
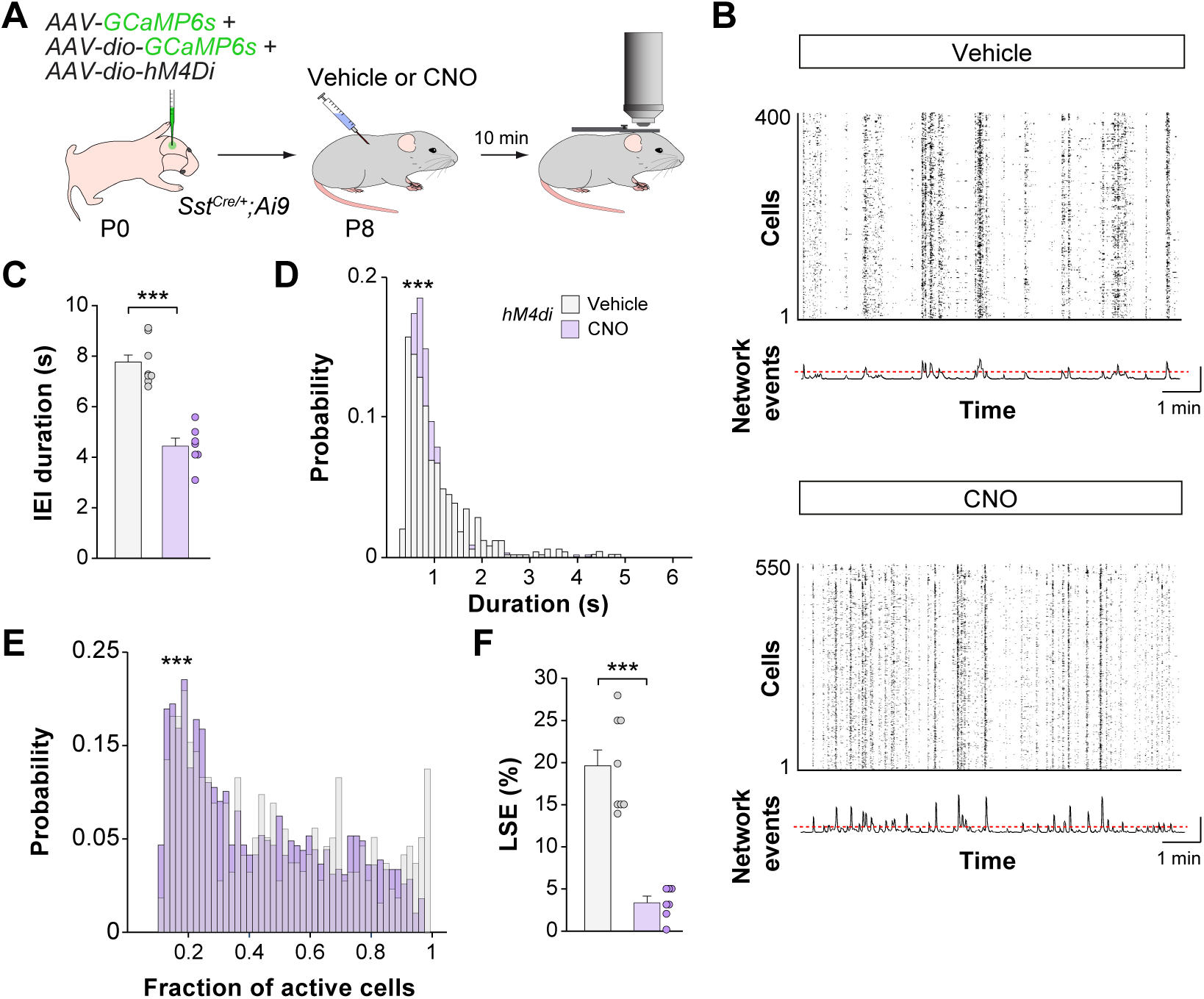
Silencing of SST+ interneurons restrict the integration of cells into High-synchronous network events. (A) Schematic representation of the experimental paradigm. We imaged 8 FOVs from 5 vehicle-injected mice and 7 FOVs from 4 CNO-injected mice. (B) Raster plots illustrating calcium traces from vehicle- and CNO-injected mice. The cumulative histogram of network events indicates the sum of active contours in the raster plots over time. The red dotted line indicates the sum of spikes considered a network event based on a statistical threshold (*p* < 0.01). (C) Quantification of inter-event intervals (IEI) duration in vehicle- and CNO-injected mice. *t*-test: *\*\*\*p* < 0.0001. (D) Frequency distribution of events length in vehicle- and CNO-injected mice. Kolmogorov-Smirnoff test: \*\*\**p* < 0.0001. (E) Distribution of probability of active cell participation in spontaneous events in vehicle- and CNO-injected mice. Kolmogorov-Smirnoff test: \*\*\**p* < 0.0001. (F) Quantification of the fraction of Large Synchronous Events (LSEs) in vehicle- and CNO-injected mice. *t*-test: \*\*\**p* < 0.0001. Data is presented as mean ± s.e.m. Each dot in (C) and (F) represents a FOV.

We next examined the impact of modulating the activity of prospective PV+ interneurons in the generation of synchronous patterns in the early postnatal cortex. To this end, we injected S1BF of P0 *Tac1^Cre/+^;Ai9* pups with a cocktail of pan-neuronal and Cre-dependent AAVs expressing GCaMP6s and Cre-dependent hM4Di-expressing AAVs, injected these mice CNO or vehicle at P8 and monitored neuronal activity 10 mins later using two-photon calcium imaging in vivo (Figure S4A). In contrast to the previous experiments, reducing the activity of prospective PV+ cells by CNO administration at P8 (Figures S3E–S3H) did not change the frequency of network events or the fraction of LSEs compared to control mice (Figures S4B–S4F). Altogether, these experiments indicate that SST+ interneurons, but not PV+ cells, shape the generation of spontaneous activity at the beginning of the second week of postnatal development.

### Whisker stimulation modulates early cortical dynamics through SST+ cells

Our previous results revealed that the activity of SST+ interneurons is required for the generation of LSEs in S1BF at the beginning of the second week of postnatal development. During this period, deep-layer SST+ interneurons receive dense thalamocortical input^26,33^, suggesting that the thalamus may influence cortical dynamics by modulating SST+ cells. To test this hypothesis, we first investigated the impact of whisker stimulation on network activity in *Sst^Cre/+^;Ai9* mice using calcium imaging in vivo during the second postnatal week. At this age, the movement of the whiskers is elicited by grooming from the dam or contact with other littermates because active whisking is still absent^41^. We mimicked this natural condition by delivering soft air puffs to the pup’s snout that activated multiple whiskers simultaneously (Figure 4A). We found that whisker stimulation does not change the frequency of events but increases the fraction of LSEs at P8 (Figures 4B and 4C). In agreement, we observed increased cell recruitment time-locked to whisker stimulation (Figure 4D). In contrast, whisker stimulation decreased the participation of cells in events at P12, including the small fraction of LSEs still present at that age (Figures S5A–S5D). This age-related effect of whisker stimulation of cortical dynamics likely reflects changes in circuit connectivity^26,33,42^ and indicates that sensory input modulates the functional organization of internal dynamics at different stages of development.

**Figure 4.**
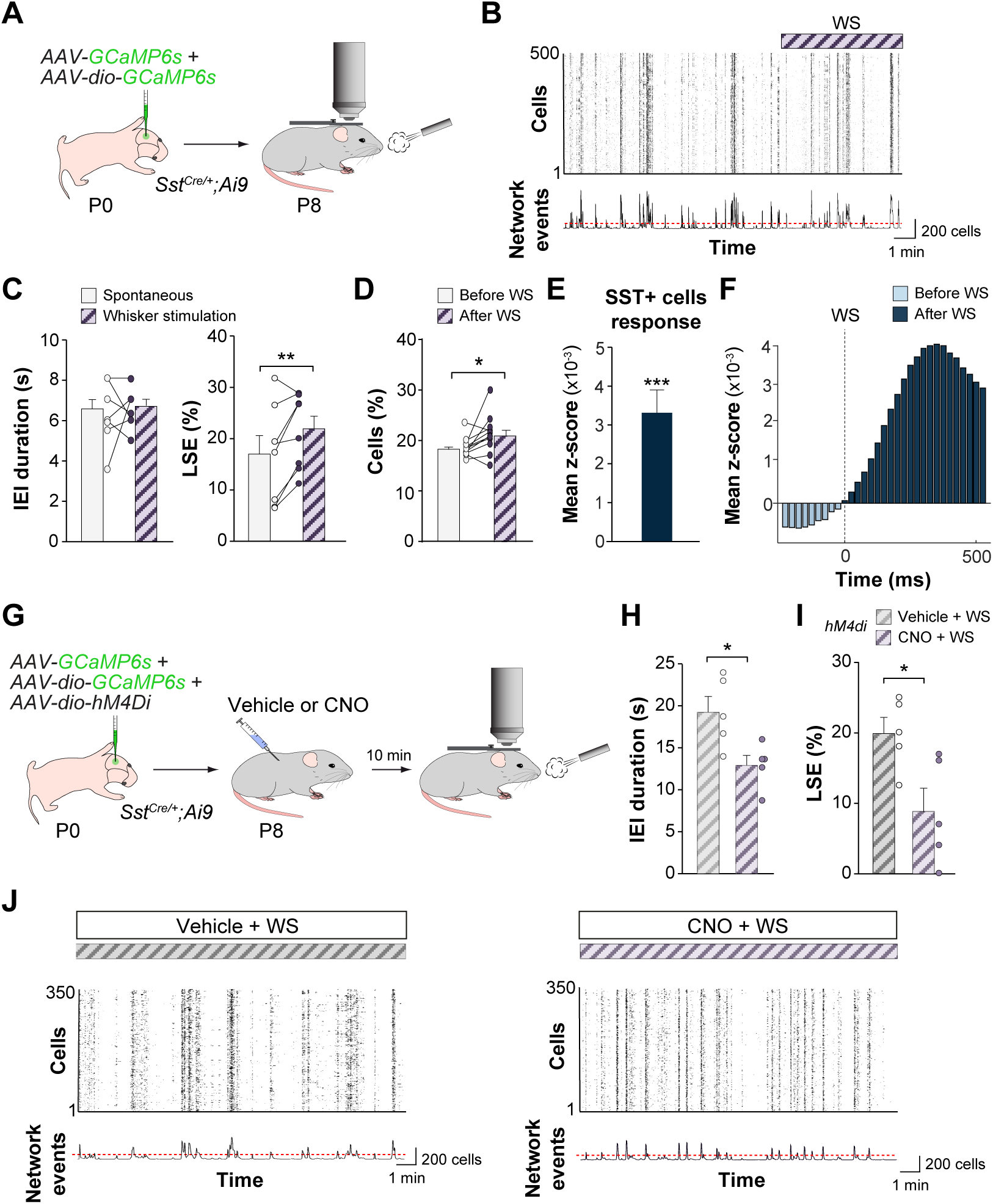
Whisker stimulation regulates cortical dynamics through SST+ interneurons. (A) Schematic representation of the experimental paradigm. We imaged 8 FOVs from 4 mice. (B) The raster plots illustrate cortical neurons’ spontaneous activity and how whisker stimulation (WS) modifies it. The cumulative histogram of network events indicates the sum of active contours in the raster plots over time. The red dotted line indicates the sum of spikes considered a network event based on a statistical threshold (*p* < 0.01). (C) Quantification of inter-event intervals (IEI) duration (left) and the fraction of Large Synchronous Events (LSEs) (right) during spontaneous activity and WS at P8. IEI, paired *t*-test: *p* = 0.78. LSEs, paired *t*-test: \*\**p* = 0.01. (D) Quantification of the fraction of active cells (%) 1s before and 1s after WS at P8. Paired *t*-test: \**p* <0.05. (E) Quantification of Mean SST+ INs response to WS at P7/8. *t*-test: \*\*\**p* <0.0001. (F) Peristimulus time histogram (PSTH) indicating changes in the z-score responses of SST+ cells before and after WS at P8. (G) Schematic representation of the experimental paradigm. We imaged 5 FOVs from 3 vehicle-injected mice and 5 FOVs from 3 CNO-injected mice. (H) Quantification of inter-event intervals (IEI) duration in vehicle- and CNO-injected mice during WS at P8. *t*-test: *\*p* < 0.05. (I) Quantification of the fraction of Large Synchronous Events (LSEs) in vehicle- and CNO-injected mice during WS at P8. *t*-test: \**p* < 0.05. (J) Raster plots illustrating calcium traces from vehicle- and CNO-injected mice during WS at P8. The cumulative histogram of network events indicates the sum of active contours in the raster plots over time. The red dotted line indicates the sum of spikes considered a network event based on a statistical threshold (*p* < 0.01). Data is presented as mean ± s.e.m. Each dot in (C) and (G) represents a FOV.

We next examined the specific response of SST+ interneurons to whisker stimulation in *Sst^Cre/+^;Ai9* mice by isolating the calcium transients of tdTomato+ cells from these experiments. We found that SST+ cells have a time-locked response to whisker stimulation at P8 and P12 (Figures 4E, 4F, and S5E–S5F). However, on average, SST+ cells respond much faster to whisker stimulation at P8 than at P12 (Figure S5G). The responses observed at P12 are reminiscent of those described in the adult cortex following whisker stimulation, where SST+ interneurons respond with a longer latency than pyramidal cells and PV+ interneurons^43^.

The previous results suggested that SST+ cells may play a more active role in modulating cortical dynamics at P8 than at P12. To test this idea, we conducted experiments in which we injected S1BF of P0 *Sst^Cre/+^;Ai9* pups with a cocktail of pan-neuronal and Cre-dependent AAVs expressing GCaMP6s and Cre-dependent hM4Di-expressing AAVs, injected these mice CNO or vehicle at P8 and monitored neuronal activity 10 mins later using two-photon calcium imaging in vivo (Figure 4G). We found that inhibiting SST+ interneurons during whisker stimulation increases the frequency of events and reduces LSE (Figures 4H–4J), suggesting that SST+ cells are indeed essential mediators of the effect of sensory stimulation on cortical dynamics at the beginning of the second week of postnatal development. In agreement with their limited participation in network events at P8 (Figure 2F), inhibiting prospective PV+ interneurons during whisker stimulation does not impact cortical dynamics at this age (Figure S6). Altogether, our results revealed that whisker stimulation promotes the highly correlated pattern of activity that characterizes S1BF at the beginning of the second week of postnatal development and that this effect is, at least in part, mediated by SST+ interneurons.

### SST+ cells regulate the decorrelation of spontaneous activity

Our previous results were based on the acute inhibition of SST+ cells. To extend these observations, we interfered with the activity of SST+ interneurons during 48 h at the beginning of the second week of postnatal development and examined the impact of this manipulation on the decorrelation of neural activity over the next few days. In a series of experiments, we injected S1BF of P0 *Sst^Cre/+^;Ai9* pups with a cocktail of pan-neuronal and Cre-dependent AAVs expressing GCaMP6s and Cre-dependent hM4Di-expressing AAVs to reduce the activity of SST+ interneurons. We injected these mice CNO or vehicle twice daily from P8 to P10 and monitored neuronal activity daily from P9 until P12 using two-photon calcium imaging in vivo (Figure 5A). We found that reducing the activity of SST+ interneurons at the beginning of the second week of postnatal development accelerates the decorrelation between cortical neurons and decreases the fraction of LSE in S1BF, which in both cases acquired values like those only typically observed at P11/12 (Figures 5B–5D). These experiments reinforced the idea that SST+ cells play a critical role in modulating cortical dynamics at the beginning of the second week of postnatal development.

**Figure 5.**
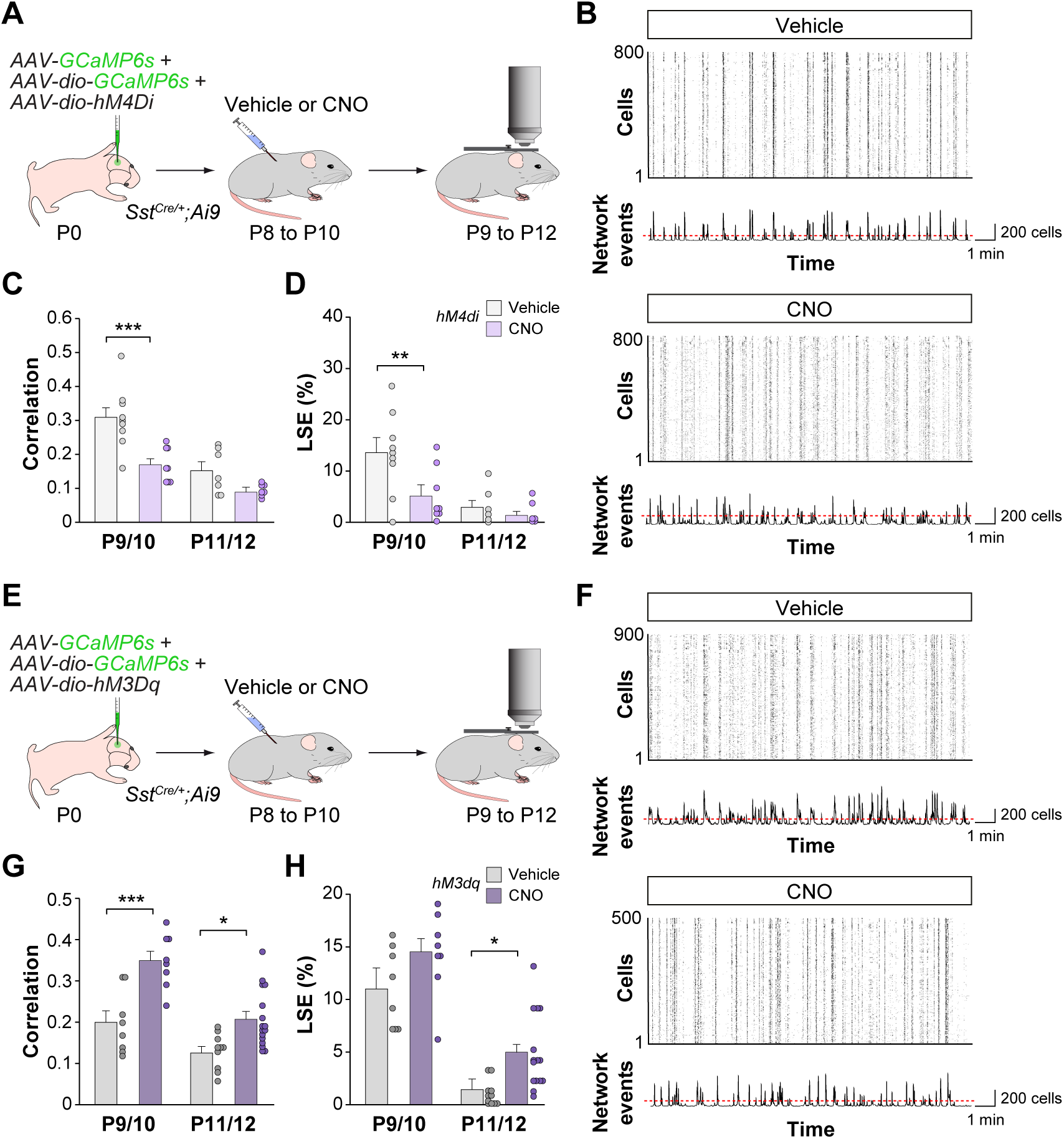
Modulation of SST+ interneurons regulates the decorrelation of spontaneous activity. (A) Schematic representation of the experimental paradigm. We imaged 9 (P9/10) and 7 (P11/12) FOVs from 3 vehicle-injected mice and 8 (P9/10) and 7 (P11/12) FOVs from 3 CNO-injected mice. (B) Raster plots illustrating calcium traces from vehicle- and CNO-injected mice. The cumulative histogram of network events indicates the sum of active contours in the raster plots over time. The red dotted line indicates the sum of spikes considered a network event based on a statistical threshold (*p* < 0.01). (C) Developmental changes in the correlation degree of active cells in vehicle- and CNO-injected mice. Two-way ANOVA for administration: F(1,27) = 17.18, *p* < 0.001; for age: F(1,27) = 25.08, *p* < 0.001. Post hoc Sidak’s correction indicates differences between vehicle and CNO-injected mice at P9/10 (\*\*\**p <* 0.001). (D) Quantification of the fraction of Large Synchronous Events (LSEs) in vehicle- and CNO-injected mice during development. Two-way ANOVA for administration: F(1,27) = 6.44, *p* = 0.01; for age: F(1,27) = 13.56, *p* = 0.001. Post hoc Sidak’s correction indicates differences between vehicle and CNO-injected mice at P9/10 (\*\**p <* 0.01). (E) Schematic representation of the experimental paradigm. We imaged 8 (P9/10) and 10 (P11/12) FOVs from 4 vehicle-injected mice and 8 (P9/10) and 16 (P11/12) FOVs from 5 CNO-injected mice. (F) Raster plots illustrating calcium traces from vehicle- and CNO-injected mice. The cumulative histogram of network events indicates the sum of active contours in the raster plots over time. The red dotted line indicates the sum of spikes considered a network event based on a statistical threshold (*p* < 0.01). (G) Developmental changes in the correlation degree of active cells in vehicle- and CNO-injected mice. Two-way ANOVA for administration: F(1,38) = 24.85, *p* < 0.0001; for age: F(1,38) = 24.85, *p* < 0.0001. Post hoc Sidak’s correction indicates differences between vehicle and CNO-injected mice at P9/10 (\*\*\**p <* 0.001) and at P11/12 (\**p <* 0.05). (H) Quantification of the fraction of Large Synchronous Events (LSEs) in vehicle- and CNO-injected mice during development. Two-way ANOVA for administration: F(1,38) =10.64, *p* < 0.01; for age: (1,38) =84.66, *p* < 0.0001. Post hoc Sidak’s correction indicates differences between vehicle and CNO-injected mice at P11/12 (\**p <* 0.05). Data is presented as mean ± s.e.m. Each dot in (C), (D), (G), and (H) represents a FOV.

In a different set of experiments, we injected S1BF of P0 *Sst^Cre/+^;Ai9* pups with a cocktail of pan-neuronal and Cre-dependent AAVs expressing GCaMP6s and Cre-dependent hM3Dq-expressing AAVs (Figure 5E). We then injected these mice CNO or vehicle twice daily from P8 to P10 and monitored neuronal activity daily from P9 until P12 using two-photon calcium imaging in vivo. We found that increasing the activity of SST+ interneurons at the beginning of the second week of postnatal development delays the decorrelation between cortical neurons and increases the fraction of LSE in S1BF, which remains relatively high even at P11/12 (Figures 5F–5H). Altogether, our results revealed that SST+ interneurons are critical regulators of the pace at which neural activity in cortical networks becomes decorrelated.

### SST+ cells modulate the maturation of PV+ interneurons

It has been suggested that the progressive strengthening of inhibition underlies the decorrelation of neural activity in the developing cortex^18,44^. The timing of PV+ cell maturation^24^, which coincides with the transition from highly synchronous to decorrelated activity in cortical networks, and the strength of PV+ synapses targeting the perisomatic region of pyramidal cells^45–47^, suggest that PV+ interneurons may play a critical role in this process. We hypothesized that SST+ interneurons modulate the decorrelation of spontaneous activity in developing cortical networks by regulating the timing of PV+ interneuron maturation. To test this hypothesis, we conducted experiments in which we imaged prospective PV+ interneurons while we interfered with the activity of SST+ cells. In brief, we injected S1BF of P0 *Lhx6-Cre;Sst^Flp/+^* pups with a cocktail of Cre-dependent AAVs expressing GCaMP6s and Flp-dependent AAVs expressing tdTomato and hM4Di. We then injected P9 mice with CNO or vehicle and monitored the activity of prospective PV+ interneurons (GCaMP6s+ and tdTomato-) using two-photon calcium imaging in vivo (Figure 6A). We found that inhibition of SST+ interneurons at this stage increases the mean spiking probability of PV+ cells and their participation in spontaneous events (Figure 6B and 6C). These results indicated that inhibiting SST+ cells facilitates the integration of PV+ interneurons into cortical dynamics at the beginning of the second week of postnatal development.

**Figure 6.**
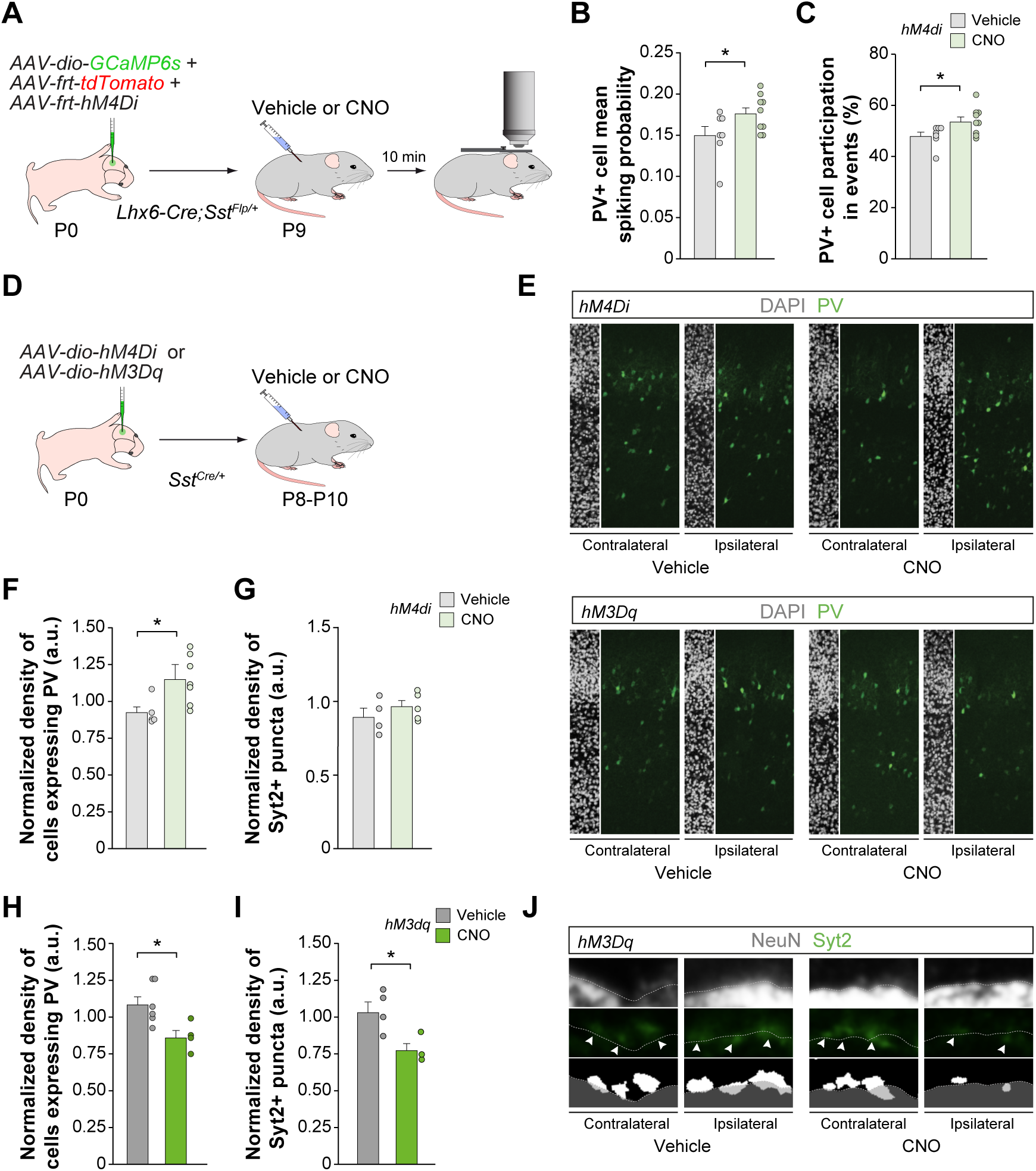
Modulation of SST+ cells regulates the maturation of PV+ interneurons and their participation in spontaneous events. (A) Schematic representation of the experimental paradigm. We imaged 7 FOVs from 4 vehicle-injected mice and 9 FOVs from 4 CNO-injected mice. (B) Quantification of the mean spiking probability of prospective PV+ cells in vehicle- and CNO-injected mice. *t*-test: *\*p* < 0.05. (C) Quantification of prospective PV+ cells participation in spontaneous events in vehicle- and CNO-injected mice. *t*-test: *\*p* < 0.05. (D) Schematic representation of the experimental paradigm. (E) Coronal sections illustrating the expression of PV in the S1BF of P10 *Sst^Cre/+^* mice injected with hM4Di or hM3Dq viruses and treated with vehicle or CNO. The ipsilateral and contralateral sides of hM4Di or hM3Dq injection are shown. (F and H) Quantification of the normalized density of cells with detectable levels of PV in the S1BF of P10 *Sst^Cre/+^* mice injected with hM4Di or hM3Dq viruses and treated with vehicle or CNO. The density of PV+ cell from the side ipsilateral to the injection was normalized by cell densities from the corresponding contralateral side to account for the brain-to-brain variability in the maturation of PV+ interneurons. hM4Di: 5 vehicle- and 7 CNO injected mice; *t*-test: \**p* < 0.05. hM3Dq: 6 vehicle- and 4 CNO injected mice; *t*-test: \**p* < 0.05. (G and I) Quantification of the normalized density of Syt2+ puncta in the S1BF of P10 *Sst^Cre/+^* mice injected with hM4Di or hM3Dq viruses and treated with vehicle or CNO. The density of Syt2+ punta from the side ipsilateral to the injection was normalized by cell densities from the corresponding contralateral side to account for the brain-to-brain variability in the maturation of PV+ interneurons. hM4Di: 4 vehicle- and 5 CNO injected mice; *t*-test: *p* = 0.39. hM3Dq: 4 vehicle- and 3 CNO injected mice; *t*-test: \**p* < 0.05. (J) Confocal images (top) and binary images (bottom) illustrating presynaptic Syt2 punta (green) contacting NeuN+ (gray) L4 excitatory cells in the S1BF of P10 *Sst^Cre/+^* mice injected with hM3Dq viruses and treated with vehicle or CNO. The ipsilateral and contralateral sides of hM3Dq injection are shown. Data is presented as mean ± s.e.m. Each dot in (B) and (C) represents a FOV. Each dot in (F), (G), (H), and (I) represents a mouse.

We next examined whether modifying the activity of SST+ interneurons promotes the integration of PV+ cells into functional networks by accelerating their maturation. To this end, we injected S1BF of P0 *Sst^Cre/+^*pups with Cre-dependent AAVs expressing inhibiting or activating DREADDs and injected these mice CNO or vehicle twice daily from P8 to P10 (Figure 6D). We then quantified the expression of PV across all cortical layers and the density of the presynaptic protein synaptotagmin 2 (Syt2) contacting L4 excitatory cells, two markers of the maturation of PV+ interneurons. We found that inhibiting SST+ cells between P8 and P10 significantly increases the density of interneurons expressing detectable levels of PV in S1BF at P10 (Figures 6E and 6F). This change was not associated with an increase in the total amount of prospective PV+ interneurons (Figure S7A-B), suggesting that inhibition of SST+ cells accelerates the onset of PV expression. No changes in the density of Syt2+ puncta contacting L4 pyramidal neurons were observed (Figure 6G). Conversely, activation of SST+ cells between P8 and P10 led to a decrease in both the density of interneurons expressing detectable levels of PV (Figures 6E and 6H) and the density of Syt2+ puncta contacting S1BF L4 pyramidal neurons at P10 (Figures 6I and 6J). The number of prospective PV+ interneurons was not altered in these experiments (Figure S7C and S7D), suggesting that the activation of SST+ interneurons delays the maturation of PV+ interneurons.

Altogether, our experiments indicate that the activity of SST+ interneurons at the beginning of the second week of postnatal development modulates the timing of maturation of PV+ interneurons and their functional integration into cortical circuits, a process that is critical for the decorrelation of neuronal activity in cortical networks.

## DISCUSSION

In this study, we set out to identify how specific subclasses of interneurons integrate and regulate the decorrelation of cortical dynamics during the first two weeks of postnatal development in the mouse cortex. We found that SST+ and PV+ interneurons integrate into functional cortical networks asynchronically, following a pattern that reflects their maturation. SST+ cells profoundly impact cortical networks at the beginning of the second postnatal, relaying sensory information and promoting the generation of the large synchronous events (LSEs) that characterize cortical activity during this period. Inhibition of SST+ interneurons accelerates the sparsification of cortical activity, suggesting that these cells are critical for timing this crucial milestone in the functional development of cortical networks. Our results indicate that the role of SST+ interneurons in this process is at least partially mediated by modulating the maturation of PV+ interneurons.

### Early spontaneous activity in the barrel cortex

The developing cortex exhibits unique patterns of spontaneous activity that drive the maturation of neural circuits^3,48–51^. At the beginning of the second postnatal week, synchronous activity in the barrel cortex is characterized by LSEs. These events have been postulated to modulate the strength of connections during circuit wiring^39,52^, which regulates cell survival during programmed cell death^15,53^. LSEs are intermingled with other events involving smaller groups of neurons, which become dominant towards the end of the second postnatal week when LSEs disappear. This binomial distribution of network events has also been reported in the developing visual cortex^49^, suggesting that the functional organization of spontaneous activity displays a similar temporal pattern across different sensory cortical regions.

Despite the similarities in the temporal evolution of spontaneous activity across cortical regions, significant differences exist in how sensory inputs modulate these events. While retinal waves traveling through the thalamus recruit small groups of V1 neurons^49,52^, our results indicate that whisker stimulation increases the fraction of LSE in S1BF during similar stages of development. Although these differences might be due to technical constraints, they are most likely caused by the divergent evolution of these cortical areas. For instance, the organization of layer 4 circuits diverges between V1 and S1BF in the adult cortex^54^, and these differences might already be present during early developmental stages. Moreover, visual information does not reach V1 until eye-opening toward the end of the second week of postnatal development, whereas S1BF integrates thalamic inputs conveying both active and passive touch.

### Role of SST+ interneurons in developing cortical dynamics

In the adult cortex, different types of interneurons modulate and synchronize network activity according to their unique anatomical and physiological specializations^21,55^. Therefore, it is not entirely surprising that their initial integration into cortical circuits dramatically shapes network activity. Inhibition in early cortical circuits is primarily conveyed by SST+ interneurons, which mature earlier than PV+ interneurons and act as transient relays of thalamic inputs^25,26,56^. Our results indicate that SST+ interneurons strongly modulate cortical dynamics at the beginning of the second week of postnatal development by promoting neuronal participation in LSEs. This observation is consistent with previous work suggesting that inhibition modulates network dynamics in the early postnatal cortex^57,58^. The effect of SST+ interneurons in network dynamics is likely mediated by the inhibition of pyramidal cells because reducing the activity of SST+ cells increases the frequency of network events. In other words, when SST+ interneurons are active, events are less frequent but recruit more cells. This paradoxical effect might be caused by the recurrent depolarization of many SST+ interneurons by pyramidal cells through synaptic facilitation and charge summation^43,59^, leading to an LSE following the release of inhibition by SST+ interneurons. Consistently, we observed that SST+ interneurons fire immediately after pyramidal cells at the beginning of the second week of postnatal development. This suggests that, at least for the barrel cortex, the long quiescent epochs that characterize cortical activity during this period might be caused by active inhibition rather than the absence of strong recurrent excitation^60^.

Our results indicate that the response of SST+ interneurons to whisker stimulation changes over the second week of postnatal development. At the beginning of this period, SST+ cells are the primary source of inhibition and rapidly respond to whisker stimulation to gate the activity of pyramidal cells in response to touch. By the end of the second week of postnatal development, however, the response of SST+ interneurons to whisker stimulation is significantly delayed, adopting a temporal pattern similar to that observed in the adult cortex^43^. This change might be caused by the progressive integration of PV+ interneurons into cortical circuits and adjustments in the connectivity of SST+ interneurons. Consistently, PV+ cells begin to fire in response to the activation of pyramidal cells towards the end of the second week of postnatal development, as observed in the adult cortex^61^. In parallel, thalamic inputs switch in target preference during this period, progressively changing SST+ cells for PV+ interneurons^25,26,33,42,62^. Thus, during this critical developmental period, SST+ interneurons in the barrel cortex work as a relay between the external world and internal dynamics, using sensory information to promote the maturation of developing local cortical circuits.

### Modulation of PV+ interneuron maturation by SST+ cells

The functional integration of PV+ cells has been postulated to underly the developmental strengthening of inhibition required for the decorrelation of neural activity in cortical circuits^18^. Although the maturation of PV+ interneurons is thought to be activity-dependent^63,64^, the molecular, cellular, and circuit-level mechanisms controlling this process remain largely unknown. Our results suggest that, during the second week of postnatal development, SST+ cells promote LSEs while delaying the maturation of PV+ interneurons. Previous studies have demonstrated that SST+ interneurons transiently connect with PV+ cells in the barrel cortex during this period^25,26,62^, which suggests that SST+ cells may gate the maturation of PV+ interneurons by reducing their excitability, firing, and integration into cell assemblies with pyramidal cells. This observation suggests a temporal division of labor for inhibitory cells during early postnatal development, with SST+ interneurons acting as the main inhibitory break of glutamatergic cells until PV+ cells have matured enough to take over the control of cortical dynamics towards the end of the second week of postnatal development. Our results indicate that the timing of that transition, marked by the desynchronization of cortical activity, is regulated by SST+ interneurons.

Changes in the number of SST+ and PV+ interneurons are often linked to alterations in cortical dynamics and have been associated with several neurodevelopmental conditions^65^. For example, aberrant neuronal synchrony underlies sensory perturbations in neurodevelopmental disorders, particularly autism^66^. Although it is unclear how early during development these deficits originate, delays in the desynchronization of cortical activity have been reported in several mouse models of neurodevelopmental disorders where the development of PV+ interneurons is compromised^67^. Our results indicate that SST+ and PV+ interneurons integrate into cortical circuits following a precise temporal sequence that coordinates the transition of neural activity from highly synchronous to sparse and decorrelated, a crucial milestone in the assembly of cortical networks.

## ACKNOWLEDGEMENTS

We thank E. Serafeimidou-Pouliou and Christoph Zimmer for general laboratory support, and I. Andrew for managing mouse colonies. We are also grateful to members of the Marín and Rico laboratories for stimulating discussions. This work was supported by a grant from the European Research Council (ERC-2017-AdG 787355) to O.M.

## AUTHOR CONTRIBUTIONS

L.M. and O.M. conceived the study. L.M. performed the calcium imaging experiments and conducted their analysis. M.M. performed and analyzed the experiments linked to the maturation of PV+ cells. L.M. and O.M. wrote the paper with input from M.M.

---

**Figure S1.**
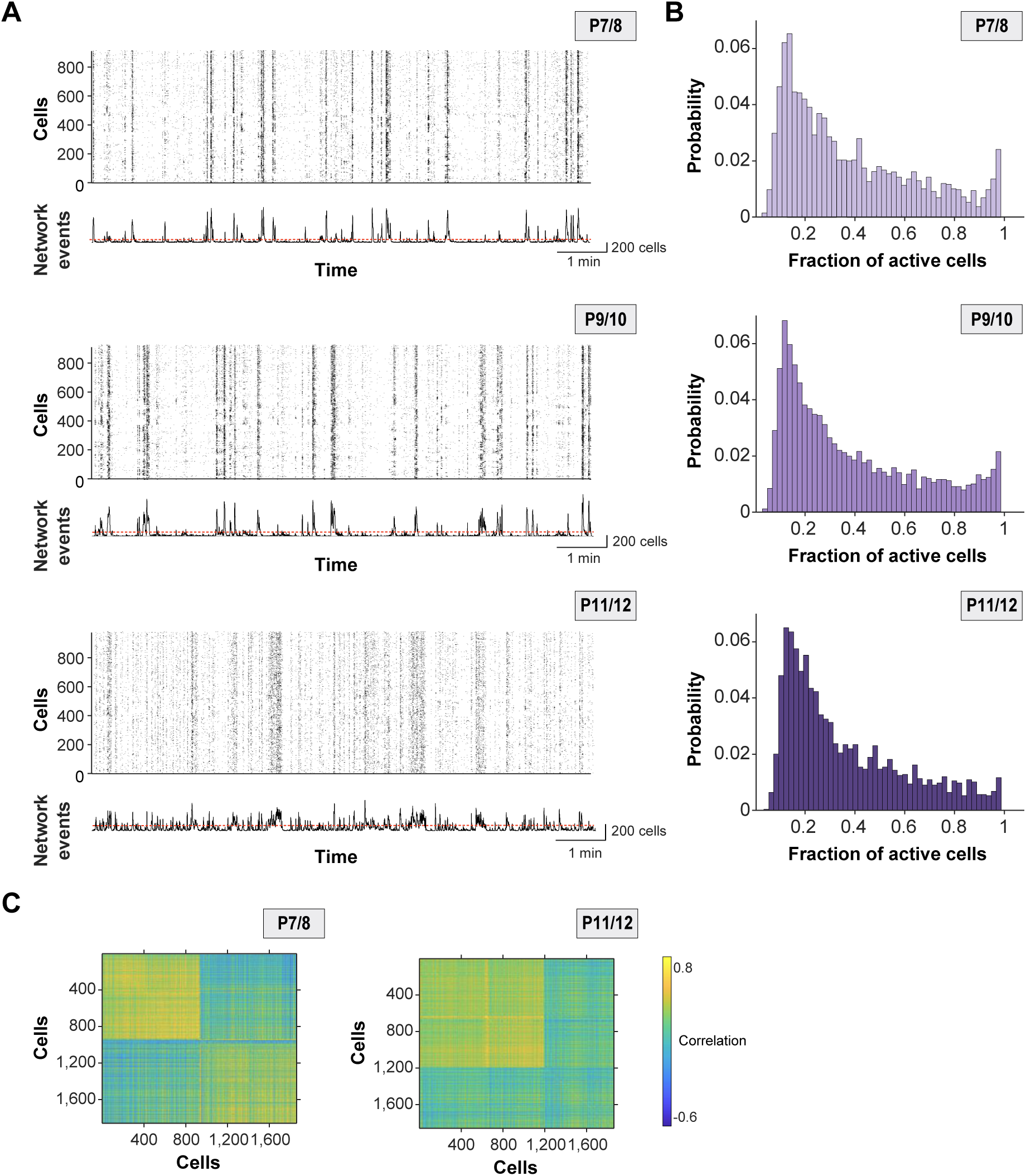
Spatiotemporal decorrelation of spontaneous activity during postnatal development. (A) Raster plots illustrating calcium events onsets during imaging sessions at P7, P9, and P11 for the same FOV. The cumulative histogram of network events indicates the sum of active contours in the raster plots over time. The red dotted line indicates the sum of spikes considered a network event based on a statistical threshold (*p* < 0.01). (B) Normalized probability of the fraction of active cells within events at different developmental stages (P7/8, P9/10, and P11/12). (C) Representative correlation matrices obtained from volumetric calcium imaging recordings of cells simultaneously imaged in L2/3 and L4/5 in S1BF at P8 and P12. These matrices correspond to the scatter plots shown in Figure 1I. Data is presented as mean ± s.e.m.

**Figure S2.**
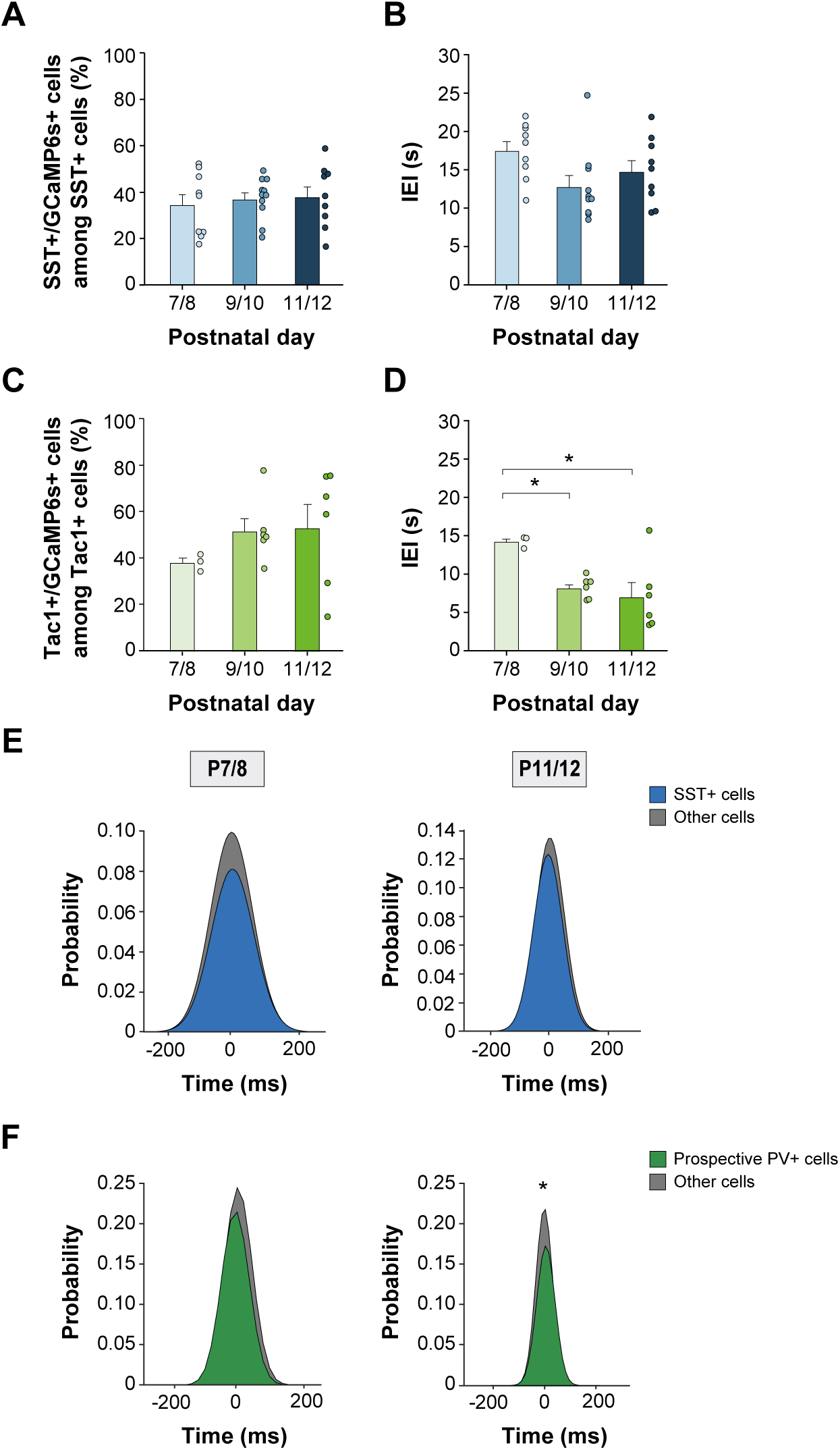
Characterization of *Sst^Cre/+^;Ai9* and *Tac1^Cre/+^;Ai9* mice. (A) Quantification of the fraction of SST+ interneurons (tdTomato+) expressing GCaMP6s in *Sst^Cre/+^;Ai9* mice as a function of age. One-way ANOVA: F(2,25) = 0.20, *p* =0.81 (9 pups at P7/8, 10 pups at P9/10, and 9 pups at P11/12). FOVs without overlapping GCaMP6s and Td-Tomato were excluded from the analysis. (B) Quantification of inter-event intervals (IEI) duration for SST+ cells in S1BF of *Sst^Cre/+^;Ai9* mice as a function of time. One-way ANOVA: F(2,25) = 2.85, *p* = 0.07. (C) Quantification of the fraction of Tac1+ interneurons (tdTomato+) expressing GCaMP6s in *Tac1^Cre/+^;Ai9* mice as a function of age. One-way ANOVA: F(2,12) = 0.70, *p* =0.51 (3 pups at P7/8, 6 pups at P9/10, and 6 pups at P11/12). FOVs without overlapping GCaMP6s and Td-Tomato were excluded from the analysis. (D) Quantification of inter-event intervals (IEI) duration for Tac1+ cells in S1BF of *Tac1^Cre/+^;Ai9* mice as a function of time. One-way ANOVA: F(2,12) = 5, *p* < 0.05. Post hoc Tukey’s multiple comparison test indicates differences between P7/8 and P9/10 (\**p* < 0.05), and between P7/8 and P11/12 (\**p* < 0.05). (E) Quantification of cell participation in spontaneous activity for SST+ cells (blue) and other active cells (grey) in *Sst^Cre/+^;Ai9* as a function of age. Shadowed distributions illustrate the probability of cells participating in events scaled to the maximum participation rate at P7/8 or P11/12*. t*-test: *p* = 0.36 and *t*-test: *p* = 0.10, respectively. (F) Quantification of cell participation in spontaneous activity for Tac1+ cells (green) and other active cells (grey) in *Tac1^Cre/+^;Ai9* as a function of age. Shadowed distributions illustrate the probability of cells participating in events scaled to the maximum participation rate at P7/8 or P11/12. *t*-test: *p* = 0.12 and *t*-test: \**p* < 0.05, respectively. Data is presented as mean ± s.e.m. Each dot in (A), (B), (C) and (D) represents a mouse.

**Figure S3.**
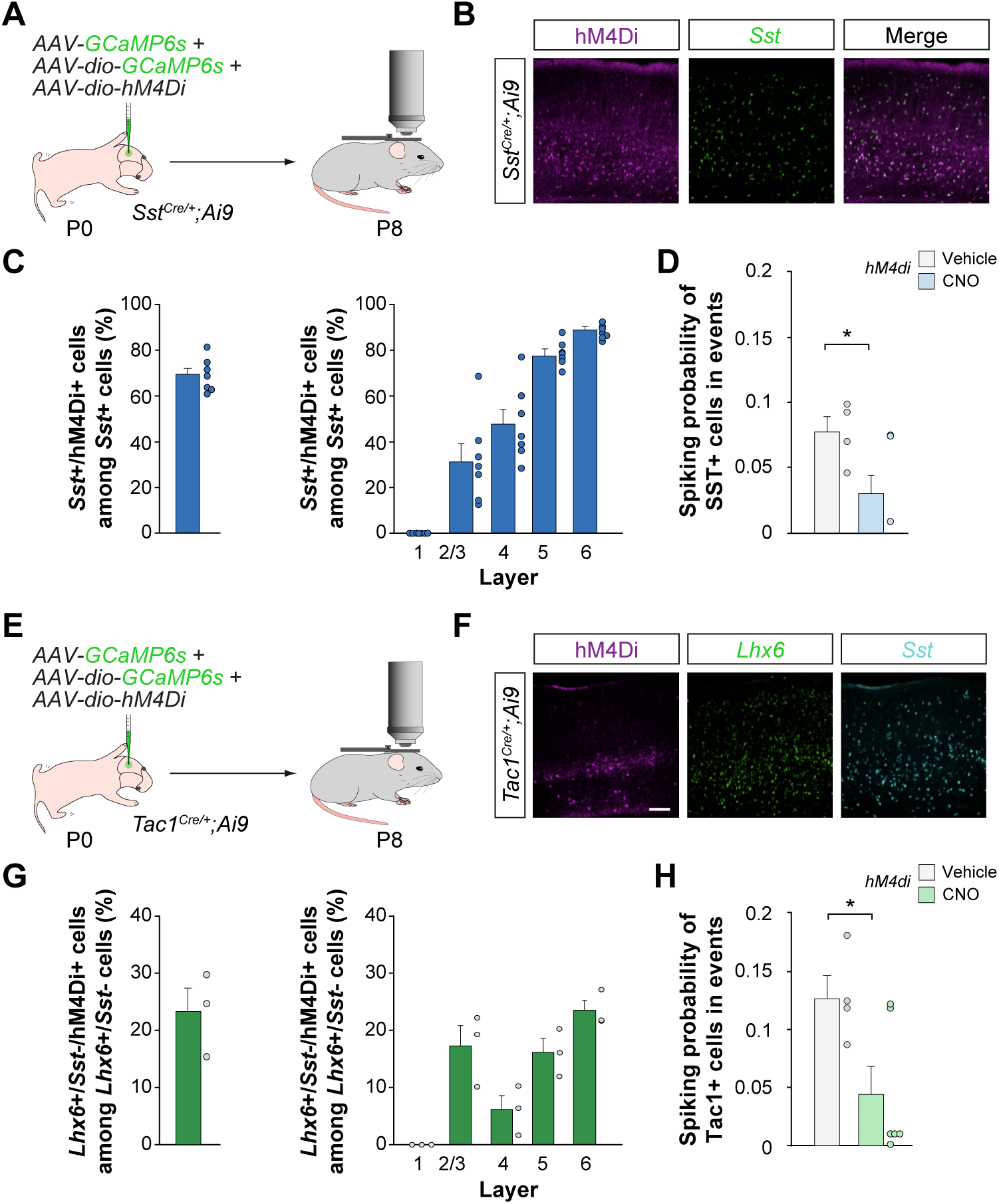
Effect of CNO in the activity of SST+ cells and prospective PV+ cells. (A) Schematic representation of the experimental paradigm. (B) Coronal sections illustrating the expression of GCaMP6s and hM4Di in S1BF of *Sst^Cre/+^;Ai9* at P8. (C) Quantification of the fraction of SST+ cells expressing hM4Di in S1BF of *Sst^Cre/+^;Ai9* at P8 (7 brains). (D) Quantification of cell spiking probability in spontaneous calcium events in vehicle- and CNO-injected *Sst^Cre/+^;Ai9* mice at P8 (4 FOVs from 3 vehicle-injected mice and 6 FOVs from 4 CNO-injected mice). *t*-test: *p* = 0.05. FOVs without overlapping GCaMP6s and Td-Tomato were excluded from the analysis (1 for vehicle-injected and 1 for CNO-injected). (E) Schematic representation of the experimental paradigm. (F) Coronal sections illustrating the expression of GCaMP6s and hM4Di in S1BF of *Tac1^Cre/+^;Ai9* at P8. (G) Quantification of the fraction of Tac1+ cells expressing hM4Di in S1BF of *Tac1^Cre/+^;Ai9* at P8 (3 brains). (H) Quantification of cell spiking probability in spontaneous calcium events in vehicle- and CNO-injected *Tac1^Cre/+^;Ai9* mice at P8 (4 FOVs from 3 vehicle-injected mice and 6 FOVs from 3 CNO-injected mice). *t*-test: *p* < 0.05. Data is presented as mean ± s.e.m. Scale bar, 100 µm (B) and (F). Each dot in (D) and (H) represents a FOV. Each dot in (C) and (G) represents a mouse.

**Figure S4.**
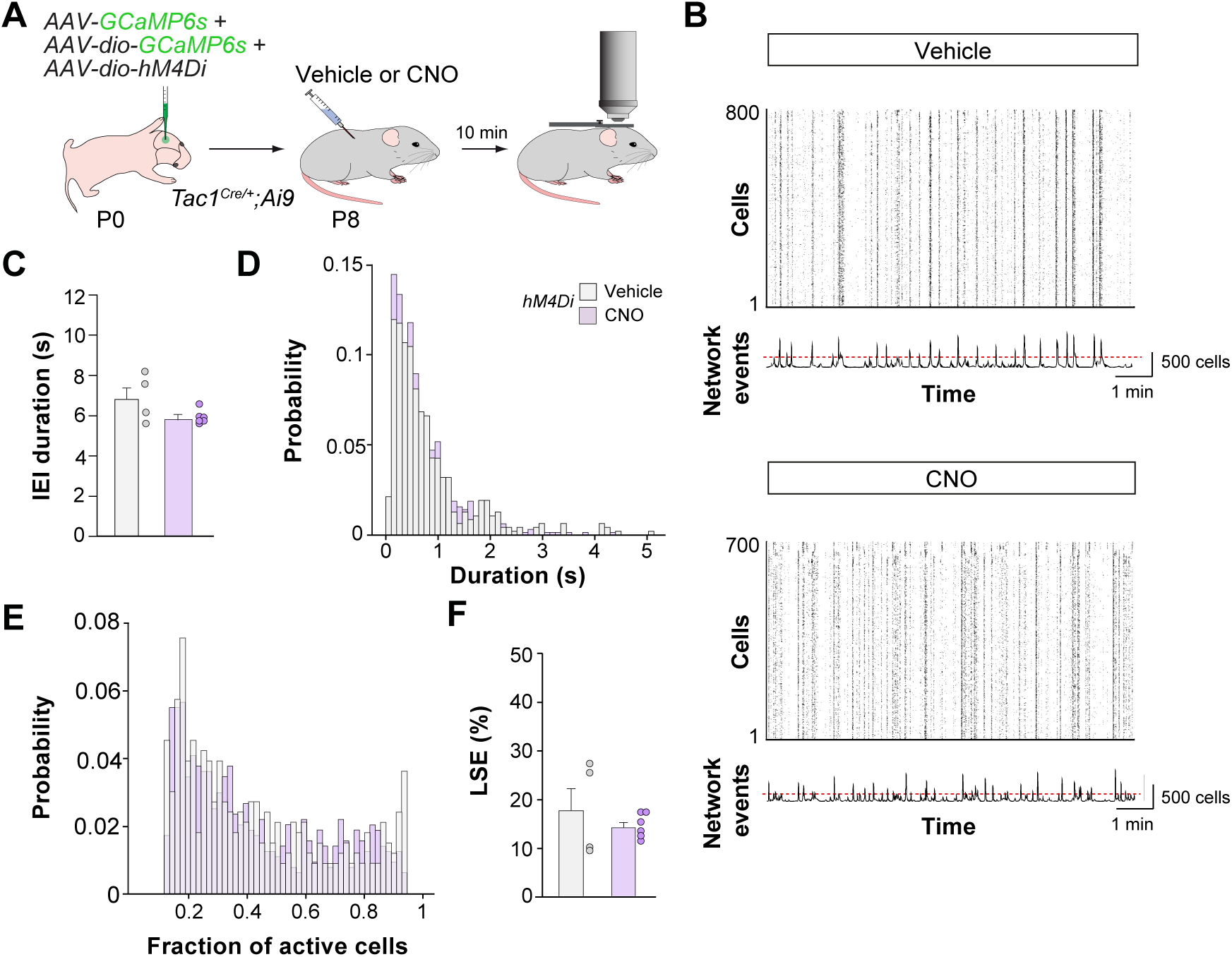
Silencing of prospective PV+ interneurons at P8 does not influence cortical dynamics. (A) Schematic representation of the experimental paradigm. We imaged 4 FOVs from 3 vehicle-injected mice and 6 FOVs from 3 CNO-injected mice. (B) Raster plots illustrating calcium traces from vehicle- and CNO-injected mice. The cumulative histogram of network events indicates the sum of active contours in the raster plots over time. The red dotted line indicates the sum of spikes considered a network event based on a statistical threshold (*p* < 0.01). (C) Quantification of inter-event intervals (IEI) duration in vehicle- and CNO-injected mice. *t*-test: *p* = 0.08. (D) Frequency distribution of events length in vehicle- and CNO-injected mice. Kolmogorov-Smirnoff test: *p* = 0.32. (E) Distribution probability of active cell participation in spontaneous events in vehicle- and CNO-injected mice. Kolmogorov-Smirnoff test: *p* = 0.06. (F) Quantification of the fraction of Large Synchronous Events (LSEs) in vehicle- and CNO-injected mice. *t*-test: *p* = 0.39. Data is presented as mean ± s.e.m. Each dot in (C) and (F) represents a FOV.

**Figure S5.**
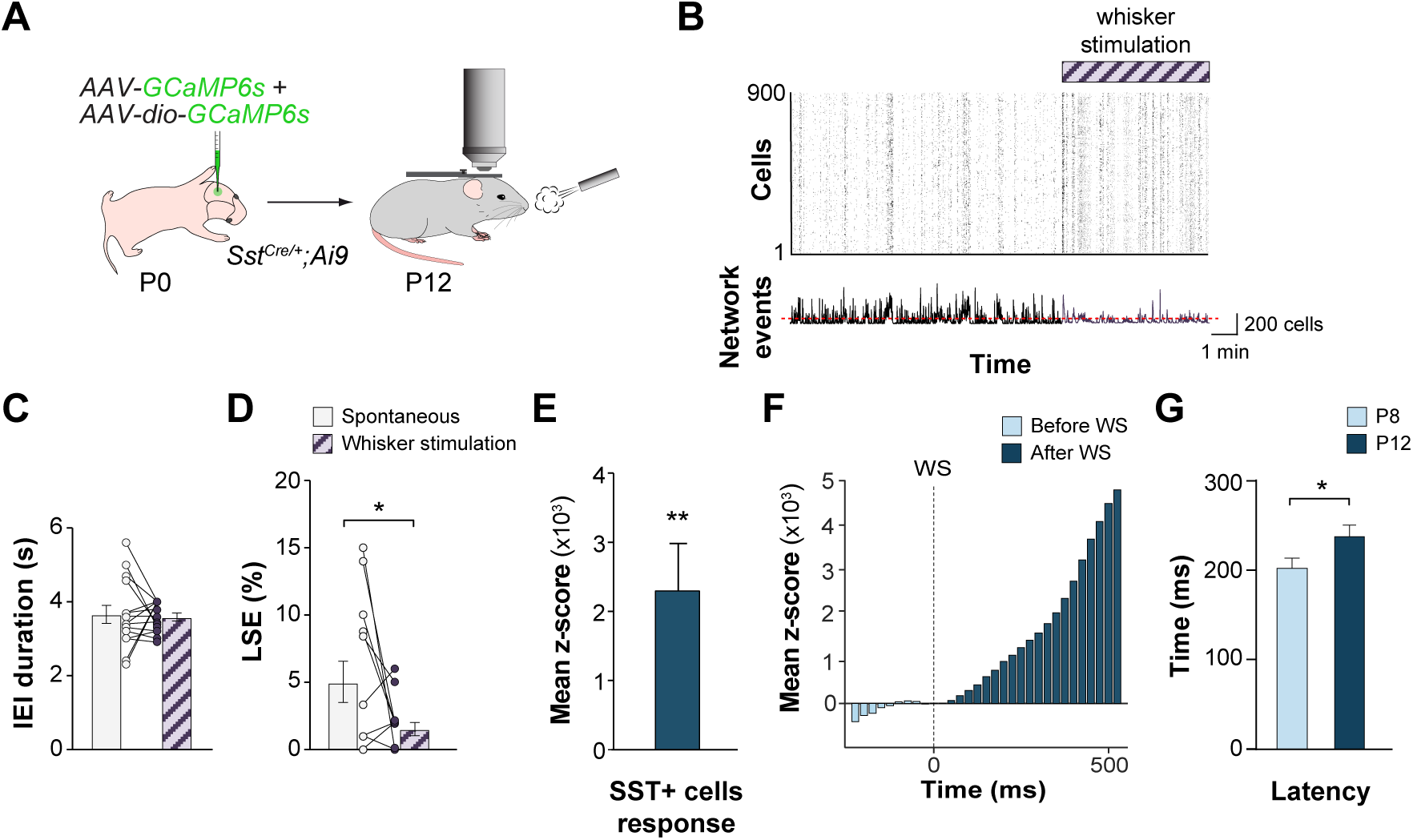
Whisker stimulation regulates cortical dynamics through SST+ interneurons. (A) Schematic representation of the experimental paradigm. We imaged 15 FOVs from 4 mice. (B) The raster plots illustrate cortical neurons’ spontaneous activity and how whisker stimulation (WS) modifies it. The cumulative histogram of network events indicates the sum of active contours in the raster plots over time. The red dotted line indicates the sum of spikes considered a network event based on a statistical threshold (*p* < 0.01). (C) Quantification of inter-event intervals (IEI) duration during spontaneous activity and WS at P12. *t*-test: *p* = 0.78. (D) Quantification of the fraction of Large Synchronous Events (LSEs) during spontaneous activity and WS at P12. Paired *t*-test: \**p* < 0.05. (E) Quantification of the mean z-score responses of SST+ cells to WS. *t*-test: \*\**p* = 0.01. (F) Peristimulus time histogram (PSTH) indicating changes in the z-score responses of SST+ cells before and after WS at P12. (G) Quantification of SST+ cells delay response (max response as seen in (F)) to WS at P8 and P12. *t*-test: * *p* < 0.05. Data is presented as mean ± s.e.m. Each dot in (C) and (D) represents a FOV.

**Figure S6.**
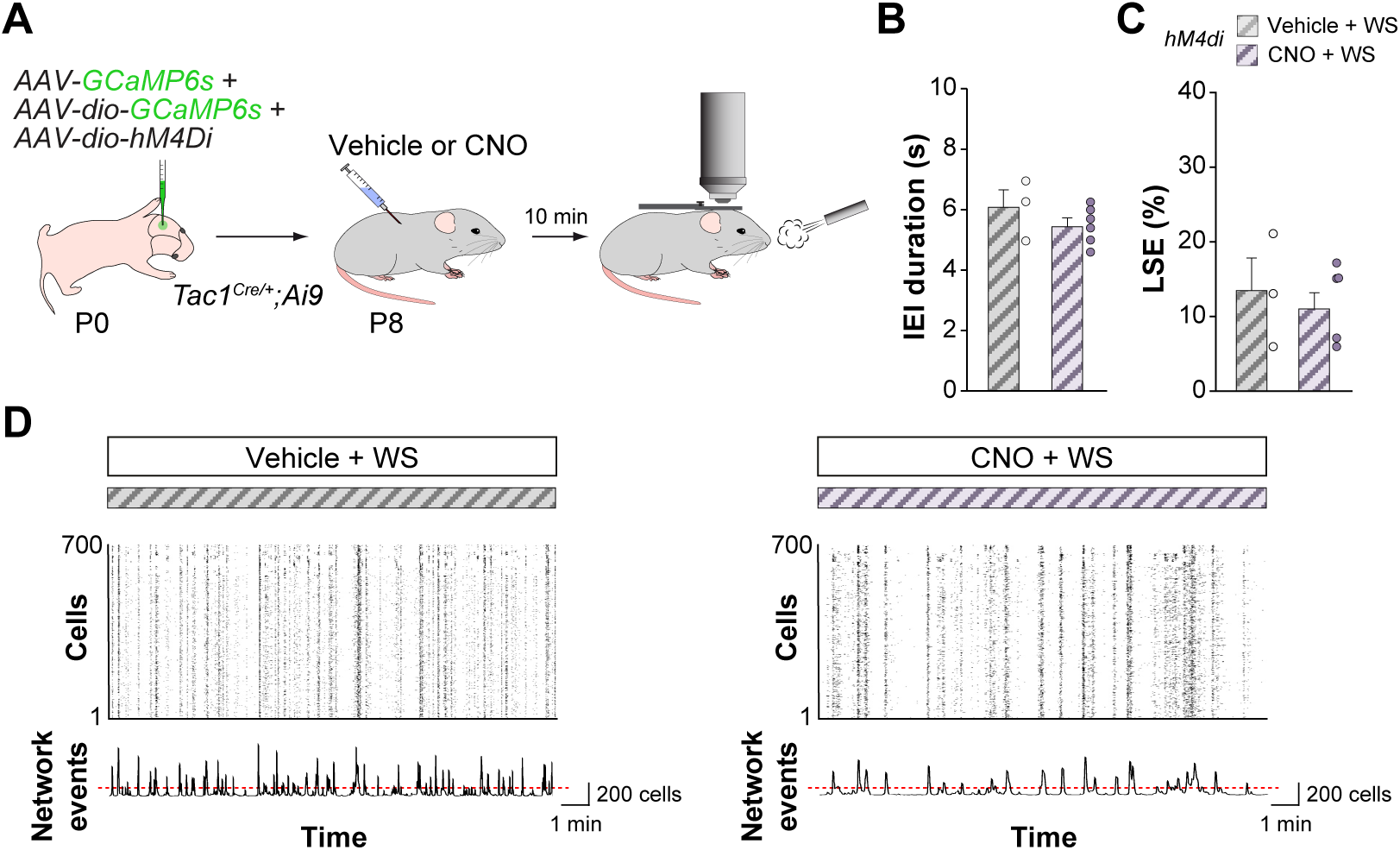
Acute modulation of prospective PV+ INs does not impact sensory-evoked responses. (A) Schematic representation of the experimental paradigm. We imaged 3 FOVs from 3 vehicle-injected mice and 6 FOVs from 3 CNO-injected mice. (B) Quantification of inter-event intervals (IEI) duration in vehicle- and CNO-injected mice during WS at P8. *t*-test: *p* = 0.29. (C) Quantification of the fraction of Large Synchronous Events (LSEs) in vehicle- and CNO-injected mice during WS at P8. *t*-test: *p* = 0.57. (D) Raster plots illustrating calcium traces from vehicle- and CNO-injected mice during WS at P8. The cumulative histogram of network events indicates the sum of active contours in the raster plots over time. The red dotted line indicates the sum of spikes considered a network event based on a statistical threshold (*p* < 0.01). Data is presented as mean ± s.e.m. Each dot in (B) and (C) represents a FOV.

**Figure S7.**
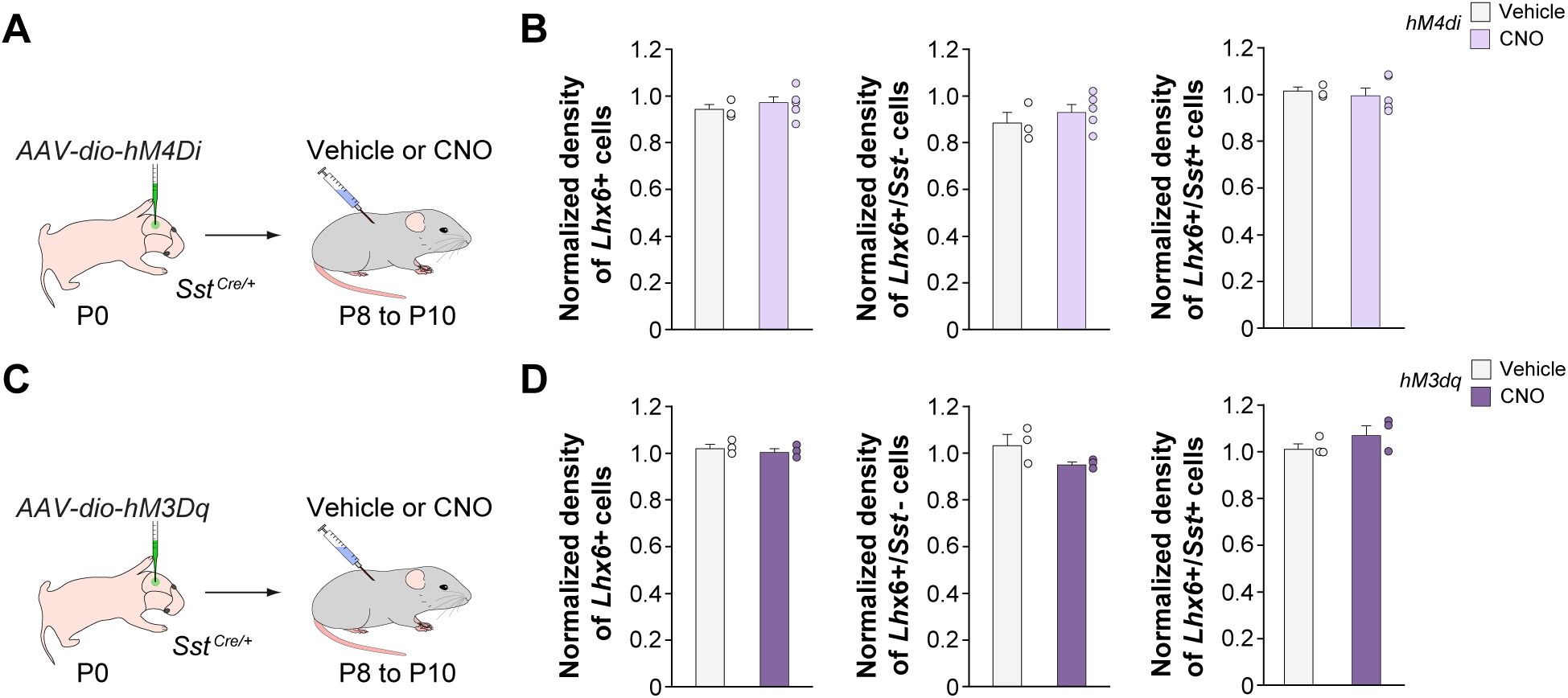
Modulation of SST+ interneurons between P8 and P10 does not alter the number of SST+ or prospective PV+ interneurons. (A) Schematic representation of the experimental paradigm. *Sst^Cre/+^* mice were injected with hM4Di virus and treated with vehicle or CNO from P8 to P10. 3 vehicle- and 5 CNO-injected pups. (B) Quantification of the normalized density of cells *Lhx6*+ in S1BF at P10. *t*-test: *p* = 0.53. Quantification of the normalized density of prospective PV+ cells (*Lhx6*+/*Sst*-) in S1BF at P10. *t*-test: *p* = 0.38. Quantification of the normalized density of SST+ cells (*Lhx6*+/*Sst*+) in the S1BF at P10. *t*-test: *p* = 0.84. Cell densities from the side ipsilateral to the injection were normalized by cell densities from the corresponding contralateral side to account for the brain-to-brain variability in the maturation of PV+ interneurons. (C) Schematic representation of the experimental paradigm. *Sst^Cre/+^* mice injected with hM3Dq virus and treated with vehicle or CNO from P8 to P10. 3 vehicle- and 3 CNO-injected pups. (D) Quantification of the normalized density of cells *Lhx6*+ in S1BF at P10. *t*-test: *p* = 0.49. Quantification of the normalized density of prospective PV+ cells (*Lhx6*+/*Sst*-) in S1BF at P10. *T*-test: *p* = 0.15. Quantification of the normalized density of SST+ cells (*Lhx6*+/*Sst*+) in S1BF at P10. *t*-test: *p* = 0.24. Cell densities from the side ipsilateral to the injection were normalized by cell densities from the corresponding contralateral side to account for the brain-to-brain variability in the maturation of PV+ interneurons. Data is presented as mean ± s.e.m. Each dot in (C) and (D) represents a mouse.

## METHODS

### Mice

All adult mice were housed in groups and kept on a 12 h light/dark cycle with ad libitum access to food and water. Only time-mated pregnant female mice were housed individually. Male and female mice were used in all experiments. For calcium imaging experiments, *Sst^Cre/+^*mice [B6N.Cg-Ssttm2.1(cre)Zjh]^68^ and *Tac1^Cre/+^* mice [B6;129S-Tac1^tm1.1(cre)Hze/J]69^ were crossed with *Ai9* mice [B6.Cg-Gt(ROSA)26Sor^tm9(CAG–tdTomato)Hze/J]70^, allowing the identification of SST+ and prospective PV+ interneurons (Tac1+) by the expression of TdTomato during the recordings. DREADDs experiments were conducted in mice obtained from crossing *Sst^Cre/+^* and *Tac1^Cre/+^* with CD1 mice. To image PV+ cell activity while modulating SST+ interneurons, we crossed *Lhx6-Cre* mice [B6;CBA-Tg(Lhx6-icre)1Kess/J]^71^ with *Sst^Flp/+^* mice [Sst^tm3.1(46lop)Zjh/J]72^. All procedures were approved by King’s College London animal welfare committees and were performed according to the project license of the UK Home Office.

### Viral production

Adeno-associated viruses (AAVs) were produced in HEK293FT cells grown on 5 (Polysciences Europe, 23966-100) or 10 15-cm diameter plates (Sigma-Aldrich, 408727), depending on whether we used linear or branched polyethylenimine (PEI) transfection, respectively, until cells reached 60% confluency. Cells were grown on DMEM (Gibco 21969-035) supplemented with 10% fetal bovine serum (FBS) (Gibco, 10500-064), 1% penicillin/streptomycin (Gibco, 15140-122), and 10 mM HEPES. AAVs were produced using PEI transfection of HEK293FT cells with a virus-specific transfer plasmid (70 µg/10 plates) and a pDP8.ape helper plasmid (300 µg/10 plates; PF478 from PlasmidFactory). The helper plasmid provided the AAV Rep and Cap functions and the Ad5 genes (VA RNAs, E2A, and E4). The DNA and PEI were mixed in a 1:4 ratio in uncomplemented DMEM and left at room temperature for 25 mins to form the DNA-PEI complex. The transfection solution was added to each plate and incubated for 72 hr. at 37 °C in 5% CO_2_. The transfected cells were then scraped off the plates and pelleted. The cell pellet was lysed in buffer containing 50 mM Tris-Cl, 150 mM NaCl, 2 mM MgCl_2_, and 0.5% Sodium deoxycholate and incubated with 100 U/ml Benzonase nuclease (Sigma, E1014 25KU) for 1 hr. to dissociate particles from membranes. The particles were cleared by centrifugation, and the clear supernatant was filtered through 0.8 µm (Merck Millipore, SLAA033SS) and 0.45 µm (Merck Millipore, SLHA 033SS) filters. The viral suspension was loaded on a discontinuous iodixanol gradient using four layers of different iodixanol concentrations^73^ of 15%, 25%, 40%, and 58% in Quick-seal polyallomer tubes (Beckman Coulter, 342414) and spun in a Vti-50 rotor at 50.000 rpm for 1 hr. 15 min at 12 °C in an Optima L-100 XP Beckman Coulter ultracentrifuge to remove any remaining contaminants. After completion of the centrifugation, 5 ml was withdrawn from the 40-58% interface using a G20 needle. The recovered virus fraction was purified by first passing through a 100 kDa molecular weight cutoff (MWCO) centrifugal filter (Sartorius, VIVASPIN VS2041) and then through an Amicon Ultra 2 ml Centrifugal filter (Millipore, UFC210024). Storage buffer (350 mM NaCl and 5% Sorbitol in PBS) was added to the purified virus, and 5 µl aliquots were stored at -80 °C.

### Viral injection

Pups were injected at birth (P0) with a viral cocktail containing *pAAV1-syn-GcaMP6s-WPRE-SV40* (Addgene, 100843-AAV1) and *pAAV1-syn-GcaMP6s-Flex WPRE-SV40* (Addgene, #100845-AAV1). In DREADDs experiments, pups were additionally injected with *pAAV8-hSyn-DiO-hM4D(Gi)-mCherry* (Addgene, 50459-AAV8) or *pAAV8-hSyn-DIO-hM3D(Gq)-mCherry* (Addgene, 44361-AAV8)*. Lhx6-Cre;Sst^Flp/+^* mice were injected with *pAAV1-Ef1a-fDiO-tdTomato* (Addgene, 128434-AAV1), *pAAV8-hSyn-fDIO-hM4D(Gi)-mCherry* (this study), and *pAAV-hSyn-FLExFRT-mGFP-2A-Synaptophysin-mRuby,* a gift from Liqun Luo^74^. Mice were separated from the dam and anesthetized with 1.5% isoflurane via a nose cone and placed in a stereotaxic frame containing a heating pad. Glass micropipettes with a tip diameter of 30-45 µm were attached to a Nanoject (Drummond) injector. Pups were injected with 600 nl of virus diluted in PBS, 150 µm below the surface, and delivered in two locations in S1BF (RC: 0.5mm; ML:1.85mm) at an injection rate of 10 nl/s. The location of the injection sites was confirmed post-mortem. Pups were allowed to recover on a heating pad at 37 °C and then returned to the dam.

### CNO administration

Clozapine-N-Oxide (CNO, Tocris, 4936) was dissolved in 5% dimethyl sulfoxide (Sigma) and then diluted with 0.9% saline to 0.1 mg/ml or 0.5 mg/ml. Pups were injected with vehicle (0.05% DMSO) or CNO (0.1g/ml) subcutaneously twice daily for three days (from P8 to P10) or acutely at P8, P9, or P12. Vehicle and CNO treatments were randomly assigned among littermates.

### Cranial windows

Cranial window surgeries were performed as described before^16^. In brief, betadine and lidocaine were applied to the skin tissue adjacent to the intended incision 20 min before starting the surgical procedures. Mice aged P6 or P7 were anesthetized with 1.5% isoflurane via a nose cone and placed in a stereotaxic frame containing a heating pad with bedding. All surgical instruments were sterilized. After skin removal, a custom-made head plate containing a 5 mm diameter hole was fixed to the skull using veterinary adhesive (Vetbond, 3M). Once the head plate was fixed and stabilized, the pup was placed in a warm cotton bed, and a 3 mm diameter craniotomy was performed over S1 and covered with a glass coverslip. The glass window was then sealed with uncured Kwik-sil (World Precision Instruments) and fixed to the head plate using Vetbond. The head plate was then fixed to the skull using Super Bond (DSM Dentaire). The body temperature was monitored and maintained close to physiological values (34–37°C) during the surgical process using a heating pad. Pups were allowed to recover for 60 min in cotton bedding and a heating pad following the procedure.

### Two-photon calcium imaging

Animals were imaged from P7 to P12 or P9 to P12 for about 1-2 h each depending on the number of FOVs. Acute experiments lasted 40 min in total. Movies were 18,000 frames at 512 x 512 pixels. Each movie lasted approximately 10 min. Imaging was performed with a single-beam multiphoton-pulsed resonant laser scanning system coupled to a microscope (Scientifica Multiphoton Resonant System). We tuned a Ti-sapphire excitation laser (Chameleon Ultra II, Coherent) at 920 nm to image GcaMP6-expressing cells. Images were acquired through a GaSP photomultiplier (H7422-40, Hamamatsu) using a 16X immersion objective (NIKON, NA 0.8). Using SciScan software (Scientifica), the fluorescence signal from a 600 µm^2^ field of view was acquired at 30 Hz with an average excitation power between 40 and 50 Mw. Movies from non-overlapping FOVs were performed in the same pup and used for the analysis.

### Whisker Stimulation

Whisker stimulation was performed by delivering brief air puffs to the snout of the pups on the contralateral side of the imaged hemisphere. Puffs were randomly assigned using a custom-made MATLAB algorithm controlling an air valvule connected to a 1 mm diameter plastic tube placed perpendicularly ∼20 mm in front of the whiskers. Air puffs lasted for 20 ms. The air puff pressure was adjusted to avoid a startle response.

### Immunohistochemistry

Mice were anesthetized with an overdose of sodium pentobarbital and transcardially perfused with saline, followed by 4% paraformaldehyde (PFA). Brains were postfixed 2h at 4°C, cryoprotected in 15% sucrose followed by 30% sucrose, and sectioned frozen on a sliding microtome at 40 µm. All primary and secondary antibodies were diluted in PBS containing 0.25% Triton X-100 and 2% BSA. The following antibodies were used: goat anti-mCherry (Antibodies Online, 1:500), rabbit anti-parvalbumin (1:5000, Swant), rabbit anti-NeuN (1:500, Millipore), mouse anti-synaptotagmin-2 (1:250, ZFIN). We used Alexa Fluor-conjugated secondary antibodies (Invitrogen, 1:400).

### Single-molecule fluorescent in situ hybridization

All single-molecule fluorescent *in situ* hybridization solutions were prepared in RNAse-free water and PBS. Mice were perfused as described above, and brains were postfixed overnight at 4°C, cryoprotected in 15% sucrose followed by 30% sucrose, and sectioned frozen on a sliding microtome at 30 µm. Sections were mounted on RNAse-free SuperFrost Plus slides (Thermofisher) and probed against target RNAs using the RNAscope Multiplex Fluorescent Assay v2 according to the manufacturer’s protocol (ACDbio #323110). The following probes were used: Mm-*Lhx6*-C2 (#422791-C2) and Mm-*Sst*-C3 (#404631-C3). The RNAscope protocol was then followed by immunohistochemistry against mCherry (goat anti-mCherry, Antibodies Online, 1:500) to stain cells infected with adeno-associated viruses expressing mCherry. Mounted sections were washed with PBS and blocked for 30 min in PBS containing 10% horse serum and 2% BSA. The sections were then incubated overnight at 4°C with the primary antibody. The next day, the sections were washed 3 times in PBS and incubated with a secondary antibody for 2h at room temperature. Mounted sections were finally counterstained with DAPI (5 µM) before drying and applying Mowiol-Dabco mounting medium (Sigma). Primary and secondary antibodies were diluted in PBS with 10% horse serum and 2% BSA.

## QUANTIFICATION AND STATISTICAL ANALYSIS

### Movement correction, deconvolution, and Ca2+ events detection

Movies obtained from in vivo imaging sessions were processed using the Suite2p toolbox^75^. In brief, the Suite2p pipeline was used for the registration (motion correction), detection of the region of interest (ROI), classification, and neuropil correction. Rigid and non-rigid (in blocks of 128A×128 pixels) options were used for the registration as previously described^75^. Cells and ROIs were then detected using clusters of correlated pixels with a low-dimensional data decomposition to accelerate processing. The number of ROIs was determined automatically by a threshold on pixel correlation. ROIs were then classified as somatic or non-somatic using a classifier trained on a set of human-curated ROIs. Only somatic ROIs (i.e., cells) were included in the analysis of calcium data. Manual curation was then used to correct for misclassification of ROIs when necessary and added to the somatic group. Neuropil correction was performed by subtracting the surrounding neuropil signal from each cell scaled by a factor of 0.7. All pixels attributed to cells were excluded from the neuropil trace.

Spike deconvolution was then performed using CASCADE, a deep learning-based tool to infer spike traces from calcium imaging data^30^. The deconvolved spike traces were then thresholded by their 99th percentile to obtain periods of cell activity and binarize the matrix. Calcium events were then detected by randomly reshuffling each spike vector of detected cells (somatic ROIs). In this way, we created five hundred surrogate distributions. The spikes of each frame for these distributions were computed, and the 99th percentile of the resulting surrogate “sum of spikes” vector was used as a statistical threshold. Calcium events were considered when the “sum of spikes” in frames of the recorded movie was above this threshold. To account for small drops in the events, such as a single frame falling below the threshold, we applied a median filter to the “sum of spikes” vector of the movie. Peaks above the threshold that were at least separated by 40 frames were considered calcium events. We used a window of -40 to +40 frames around the peak to establish which cells participate in calcium events. The spiking probability of cells was calculated based on the mean spike rate of each cell within the duration of each detected calcium event.

### Cell participation in events

Cell participation in calcium events was considered when a cell was firing at least once within the length of the event. The percentage of cell participation in calcium events was then calculated by obtaining the fraction of events in which a cell was firing. Cell participation was calculated for TdTomato- and TdTomato+ cells. TdTomato + cells were manually detected by overlapping the cell contour map of active cells (somatic ROIs) previously detected using Suite2p and a z-projection of 200-300 frames containing red cells recorded after each imaged FOV. Inter Event Intervals (IEI) were then calculated by dividing the length of the calcium movie by the number of detected calcium events. The mean and standard deviation of cell participation in events were calculated based on how often a cell would be active during a detected event. The mean and the standard deviation were then calculated, obtaining the pooled average of the cells within all events.

### Correlation coefficient

To calculate changes in the correlation of cell activity during postnatal development, we obtained the Mean Pearson correlation coefficient among the cells’ deconvolved traces.

### Probability density distribution

The density distribution of cell participation in events at different postnatal stages (P7/8, P9/10, and P11/12) was obtained using a kernel smoothing function (Matlab *ksdensity* function). Based on the inferred distribution, we then detected the peaks using the Matlab *findpeaks* function.

### Detection of large synchronous events

Large synchronous events (LSE) were defined as events involving more than 80% of the active cells within the length of the event. We first determine the length of the event based on the consecutive frames containing co-spiking of active cells above the previously established threshold. In this way, the length of an event was not fixed. We then computed the cells that were active at least once during these events and obtained the fraction of LSE.

### Spike distributions

The average onset time of the first spike across all events was considered the average onset of the cell. Differences between L2/3, L4, and L5, or between SST+ cells, Tac1+ cells, and other active cells, were calculated, obtaining a group-wise probability density function of onsets that were then scaled to the maximum participation rate per group and centered around 0 (were 0 is the peak of cells participating).

### Cluster analysis

Cluster analysis was performed as described before^16,76^. In brief, cells in clusters were identified using a clustering algorithm based on events similarity according to cell participation, followed by a statistical test for cell participation in each event cluster. The calcium event similarity metric was the squared Euclidian distance between columns of the normalized covariance matrix. This similarity metric allowed for a more efficient clustering. Unsupervised clustering of calcium events was obtained by running the *k-means* algorithm on this metric with cluster numbers ranging from 2 to 19. A hundred iterations of *k-means* were run for each cluster number, and the iteration that resulted in the best-averaged *silhouette* value was kept. The *silhouette* value was computed as described previously^76^. A random distribution of average silhouette values for each cluster was calculated by reshuffling cell participation across different calcium events and applying the same algorithm. Clusters with average *silhouette* values exceeding the 95th percentile of the random case were considered statistically significant. Each cluster was then associated with a cell assembly that comprised those cells that significantly participated in the calcium events within that cluster. Cell participation in a cluster was considered statistically significant if the fraction of synchronous events in that cluster that activated the cell exceeded the 95th percentile of reshuffled data. If a cell was significantly active in more than one cluster, it was associated with the one in which it participated the most (percentage-wise). The overlap between clusters was quantified by calculating the *silhouette* value of each cell (with the normalized hamming distance between each cell pair as a dissimilarity metric). A cell was significantly involved in a single cluster if its *silhouette* value was higher than expected by chance (95th percentile after reshuffling). The percentage of correlated cells was then obtained by dividing the differences in the number of cells participating in clusters (highly correlated) divided by the total number of active cells (GCaMP6s-expressing cells).

### PSTH response and latency

To obtain peristimulus time histograms (PSTH), we first normalized the dF/F of tdTomato+ cells between 0 and 1. We then obtained the mean response across all cells and subtracted the mean value at time 0 (the time point where the whisker puff was delivered) from the averaged response. Latency response to WS stimulation was then calculated by taking the frame with the maximum mean response for each cell.

### Statistical analysis

Data are given as mean and standard error of the mean (SEM) or as median and interquartile range. Statistics and graphs were performed using Prism 9 (GraphPad) or MATLAB. No statistical methods were used to predetermine sample sizes. Sample sizes were chosen based on previous publications in the field. Statistical analysis was performed using one-way, two-way ANOVA, or Kruskal Wallis to determine the interaction and dependencies between one or two factors. We applied Tukey’s HSD or Sidak’s post hoc tests to correct for multiple comparisons when necessary. All degrees of freedom for the ANOVA and the *t*-tests are indicated in the figure legends. Kolmogorov-Smirnov was used to analyze differences in cumulative probability distributions.

### Histological image acquisition and analysis

Images were acquired with an SP8 confocal microscope (Leica TCS SP8). Samples from the same experiment were imaged and analyzed in parallel, using the same laser power, photomultiplier gain, and detection filter settings. Imaging was performed at 12-bit depth for cell density analyses using a 10X objective. Images following immunohistochemistry experiments were acquired with a 1024 x 1024 pixel resolution, while imaging of in situ hybridization experiments was performed with a 2048 x 2048 pixel resolution. Parameters of 0.75 digital zoom and 400 Hz acquisition speed were used. Six to eight sections were imaged per brain hemisphere and analyzed after in situ hybridization or immunohistochemistry on the ipsilateral and contralateral sides of injection. Manual cell counting was then performed across layers of the somatosensory cortex using the FIJI (Image J) software. Cell densities from the side ipsilateral to the injection were normalized by cell densities from the corresponding contralateral side to account for the brain-to-brain variability in the maturation of PV+ interneurons.

For Syt2 puncta density analysis, imaging was performed at 8-bit depth using a 100X objective with oil, 1024 x 1024 pixel resolution, 2.2 digital zoom, and 400Hz acquisition speed. Approximately 30 NeuN+ cells were imaged per hemisphere (ipsilateral and contralateral sides). We used a custom macro in FIJI to analyze Syt2 synaptic density, as described previously^77,78^. Background subtraction, Gaussian blurring, smoothing, and contrast enhancement were first applied in all channels. Cell somas were drawn manually based on intensity levels of NeuN staining to create a mask of the soma surface and measure its perimeter. Presynaptic boutons were detected automatically based on thresholds of intensity. The threshold for Syt2 was selected from a set of random images before quantification, and the same threshold was applied to all images from the same experiment. The “Analyze Particles” and “Watershed” tools were applied to the Syt2 channel, and a mask was generated with a minimum particle size of 0.05. Puncta were defined as presynaptic boutons when they were located outside the soma and had ≥ 0.1 μm^2^ colocalizing with the soma perimeter. The mean Syt2 densities measured in the side ipsilateral to the injection were normalized by the mean Syt2 densities in the corresponding contralateral side to account for the brain-to-brain variability in the maturation of PV interneurons.

